# The formal demography of kinship VI: Demographic stochasticity, variance, and covariance in the kinship network

**DOI:** 10.1101/2024.05.22.594706

**Authors:** Hal Caswell

**Affiliations:** Institute for Biodiversity and Ecosystem Dynamics, University of Amsterdam

## Abstract

**Background:** The matrix model for kinship networks includes many demographic processes but is deterministic, projecting expected values of age-stage distributions of kin. It provides no information on (co)variances. Because kin populations are small, demographic stochasticity is expected to create appreciable inter-individual variation.

**Objectives:** To develop a stochastic kinship model to project (co)variances of kin age-stage distributions, and functions thereof, including demographic stochasticity.

**Methods:** Kin populations are described by multitype branching processes. Means and covariances are projected using matrices that are generalizations of the deterministic model. The analysis requires only an age-specific mortality and fertility schedule. Both linear and non-linear transformations of the kin age distribution are treated as outputs accompanying the state equations.

**Results:** The stochastic model follows the same mathematical framework as the deterministic model, modified to treat initial conditions as mixture distributions. Variances in numbers of most kin are compatible with Poisson distributions. Variances for parents and ancestors are compatible with binomial distributions. Prediction intervals are provided, as are probabilities of having at least one or two kin of each type. Prevalences of conditions are treated either as fixed or random proportions. Dependency ratios and their variances are calculated for any desired group of kin types. An example compares Japan under 1947 rates (high mortality, high fertility) and 2019 rates (low mortality, low fertility).

**Contribution:** Previous versions of the kinship model have acknowledged their limitation to expected values. That limitation is now removed; means and variances are easily and quickly calculated with minimal modification of code.

## 1 Introduction

The matrix kinship model (Caswell, 2019) treats the kin, of each kind, of a focal individual (called ‘Focal’) as a population. It uses projection matrices to project the expected age distribution of those kin populations as Focal ages. Since its introduction it has been extended to include time-varying demographic rates, classification by both age and stage (e.g., parity), inclusion of male and female rates, and the loss of kin by multiple causes of death (Caswell, 2020, 2022; Caswell and Song, 2021; Caswell, Verdery, and Margolis, 2023).

As with all such projections, the results are means, or expected values of the age distributions of kin. However, the populations of the various types of kin are small, and so we expect demographic stochasticity to lead to appreciable variation around those expected values.^1^

This paper presents a stochastic version of the model that projects the means, variances, and covariances of age structures and functions of those age structures, provides prediction intervals around the expected numbers of kin, and provides new insight into prevalence of conditions and dependency ratios among kin.

As with previous generalizations of the original model, the stochastic model maintains the population projection approach, but with the survival and fertility matrices enlarged to account for covariances as well as means. As we will see, the calculations are familiar, but with care required to take advantage of the stochastic information.

**Outline**. Section 2 presents the stochastic version, based on the theory of multitype branching processes, of the matrix kinship model. Section 2.3 details the construction of the stochastic projection matrix, and Section 2.4 describes the initial conditions for the stochastic projections. Given the model, Section 3 presents some potentially interesting analyses of the kinship network, including variances, standard deviations, and prediction intervals for kin numbers. An example follows, contrasting a high mortality, high fertility population (Japan under 1947 rates) with a low mortality, low fertility population (Japan under 2019 rates). Section **??** presents two more elaborate stochastic outcomes: treating prevalences as probabilities, and calculating the covariance structure of dependency ratios.

### 1.1 Notation

The following notation is used throughout this paper. Matrices are denoted by upper case bold characters (e.g., **U**) and vectors by lower case bold characters (e.g., **a**). Vectors are column vectors by default; **x**^T^ is the transpose of **x**. The *i*th unit vector (a vector with a 1 in the *i*th location and zeros elsewhere) is **e**_*i*_. The vector **1** is a vector of ones, and the matrix **I** is the identity matrix. When necessary, subscripts are used to denote the size of a vector or matrix; e.g., **I**_*ω*_ is an identity matrix of size *ω × ω*. The diagonal operator 𝒟 (**x**) creates a diagonal matrix with **x** on the diagonal.

The symbol º denotes the Hadamard, or element-by-element product (implemented by .* in Matlab and by * in R). The symbol ⊗ denotes the Kronecker product. The vec operator stacks the columns of a *m* × *n* matrix into a *mn* × 1 column vector. The notation ‖**x** ‖ denotes the 1-norm of **x**. Matlab notation will be used to refer to rows and columns of a matrix; e.g., **F**(*i*, :) and **F**(:, *j*) refer to the *i*th row and *j*th column of the matrix **F**, respectively.

## 2 The stochastic kinship model

### 2.1 The deterministic model reviewed

We review briefly the deterministic model; derivations are given in detail in Caswell (2019). The kinship network is defined relative to a focal individual (called Focal); the network is shown in Figure 1. The letters in this diagram denote the age distribution vectors of each type of kin.

**Figure 1:**
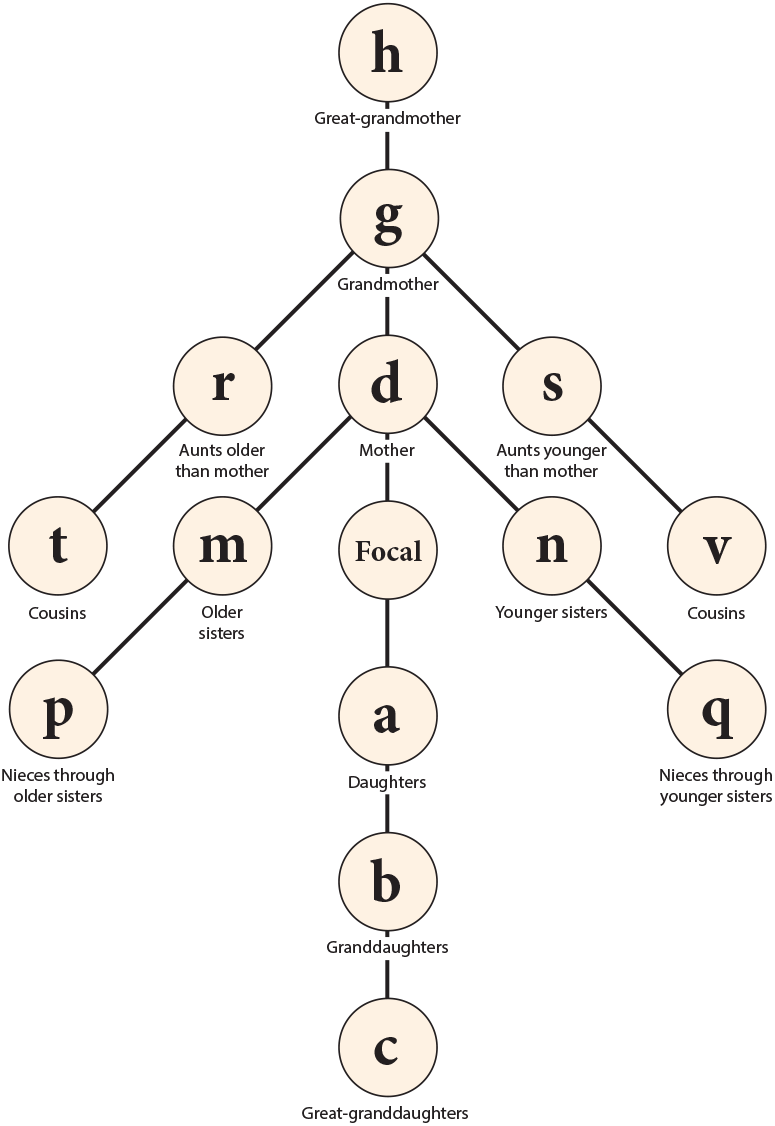
The kinship network surrounding Focal. The branches can be extended at will in all directions if desired. From Caswell (2019) under a CC-BY license.

**Figure 2:**
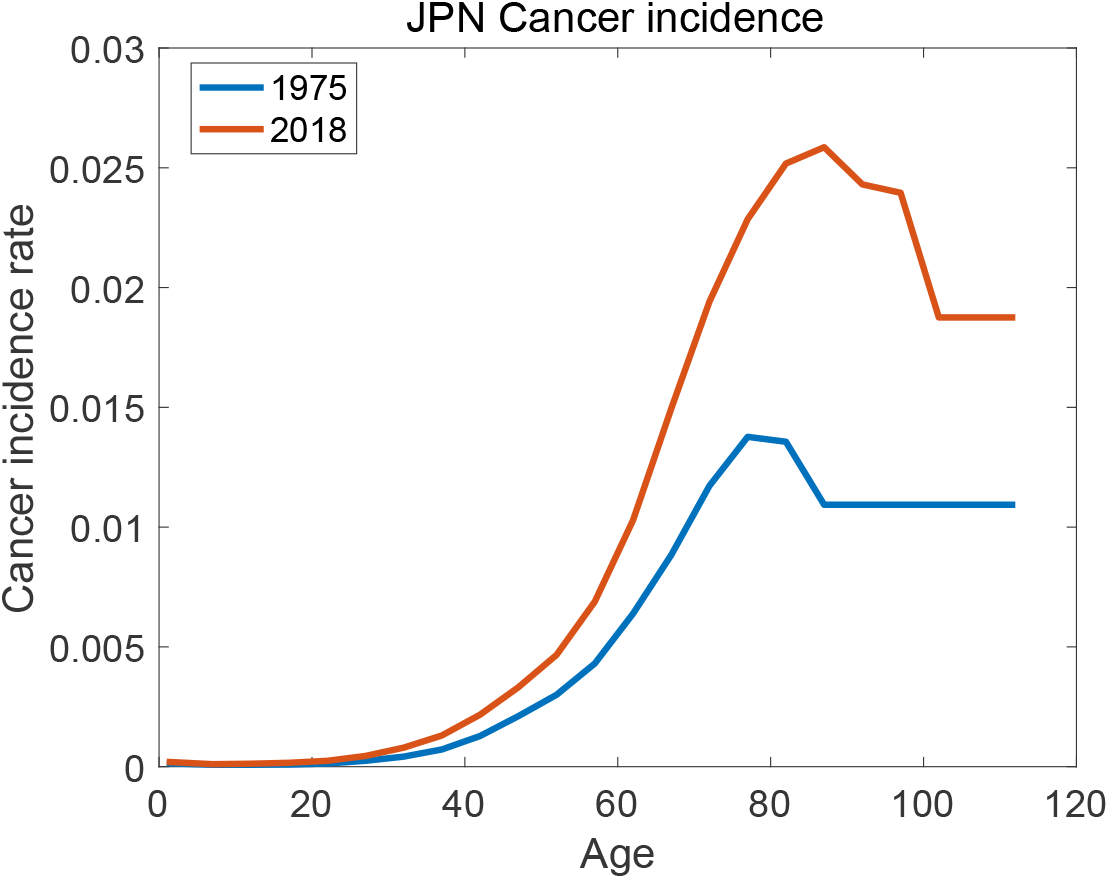
Age-specific cancer incidence rate in Japan, all causes, in 1975 and 2018. Rates have been extrapolated by being held constant after the last age reported.

Suppose the population contains *ω* age classes. Define the survival matrix **U** (size *ω* × *ω*) with survival probabilities on the subdiagonal and zeros elsewhere, and the fertility matrix **F** (also *ω* × *ω*) with age-specific fertility in the first row and zeros elsewhere. For a generic kin type **k**(*x*) at age *x* of Focal the model is

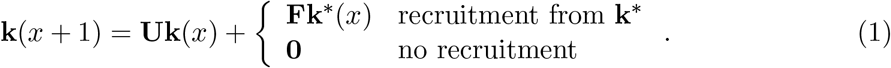

That is, the kin **k** at age *x* + 1 of Focal are the survivors of the kin at age *x* and the new recruits, if any, coming from the fertility of some other kind of kin **k**^*^ (e.g., new grandchildren come from the fertility of children).

The initial condition **k**_0_ gives the age structure of the kin at the birth of Focal. For some types of kin, it is known that the initial condition is zero (e.g., Focal has no children when she is born). For others, the kin at the birth of Focal are calculated as average over the distribution π of ages of mothers at the birth of children. The resulting calculations for each of the kin in Figure 1 are summarized in Table 1.

**Table 1:** The initial condition and recruitment term for each type of kin in the deterministic (one-sex, time-invariant) kinship model: from Caswell (2019) under a CC-BY license.

| Symbol | Kin | i.c. $\mathbf{k}_0$ | $\beta(x)$ |
| --- | --- | --- | --- |
| **a** | daughters | $\mathbf{0}$ | $\mathbf{F}\mathbf{e}_x$ |
| **b** | granddaughters | $\mathbf{0}$ | $\mathbf{F}\mathbf{a}(x)$ |
| **c** | great-granddaughters | $\mathbf{0}$ | $\mathbf{F}\mathbf{b}(x)$ |
| **d** | mothers | $\boldsymbol{\pi}$ | $\mathbf{0}$ |
| **g** | grandmothers | $\sum_i \pi_i \mathbf{d}(i)$ | $\mathbf{0}$ |
| **h** | great-grandmothers | $\sum_i \pi_i \mathbf{g}(i)$ | $\mathbf{0}$ |
| **m** | older sisters | $\sum_i \pi_i \mathbf{a}(i)$ | $\mathbf{0}$ |
| **n** | younger sisters | $\mathbf{0}$ | $\mathbf{F}\mathbf{d}(x)$ |
| **p** | nieces via older sisters | $\sum_i \pi_i \mathbf{b}(i)$ | $\mathbf{F}\mathbf{m}(x)$ |
| **q** | nieces via younger sisters | $\mathbf{0}$ | $\mathbf{F}\mathbf{n}(x)$ |
| **r** | aunts older than mother | $\sum_i \pi_i \mathbf{m}(i)$ | $\mathbf{0}$ |
| **s** | aunts younger than mother | $\sum_i \pi_i \mathbf{n}(i)$ | $\mathbf{F}\mathbf{g}(x)$ |
| **t** | cousins from aunts older than mother | $\sum_i \pi_i \mathbf{p}(i)$ | $\mathbf{F}\mathbf{r}(x)$ |
| **v** | cousins from aunts younger than mother | $\sum_i \pi_i \mathbf{q}(i)$ | $\mathbf{F}\mathbf{s}(x)$ |

### 2.2 The stochastic model: The population vector

The stochastic model projects not only the expected age distribution, but also the covariance matrix of that distribution. The augmented kin vector 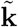 for kin of type **k** is

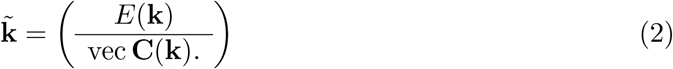

The upper block (*ω* × 1) is the expected value of the kin age distribution. The lower block (*ω*^2^ × 1) contains, in vectorized form, the covariance matrix of the age distribution. The covariance matrix of **k** is

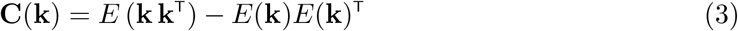

This is a good time to be reminded^2^ of some manipulations of the covariance matrix that will appear repeatedly. The covariance of a linear transformation is particularly important Appendix A.1). We will also uset the law of total covariance to partition a total covariance into within-group and between-group components (Appendix A.2).

### 2.3 The stochastic model: Projection of kin as a branching process

Branching processes are the most basic expression of demographic stochasticity. An branching process describes a population of entities that make copies of themselves. The number of copies is a random variable with a specified probability distribution. Individuals make copies independently of each other, and the population grows according to the random accumulation and loss of copies.

A multitype branching process describes a population of entities of multiple types (e.g., age classes) that produce, again independently, copies that may be of the same type or of other types. The number of copies is a random variable with specified probability distribution. Caswell (2001, Chap. 15) gives an introduction to single- and multi-type branching processes in a demographic context.

A branching process projects the probability generating function (pgf) of the population vector giving the numbers of each type. Except in special simplified cases, it is impossible to explicitly write down the pgf. However, the moments of population size are obtained from the derivatives of the pgf. The mean comes from the first derivatives, and the projection of the mean is the familiar population projection (or Leslie) matrix (Pollard, 1973). The (co)variances are obtained from the second derivative of the pgf (Harris, 1963, Eq. 4.2), and so on.

In a remarkable paper, Pollard (1966; see also Pollard 1973) used this result to arrive at a matrix that projects both means and covariances of the population structure. A detailed presentation, extended from age-classified to stage-classified populations, with an application to the orca, given in Caswell (2001, Chap. 15).

To obtain a kinship model, we must account for the linkage between kin types via recruitment. The assumption of independence of individuals means that recruitment is a random independent input to the branching process of any kin type, and hence the moments of **k**(*x*) are the sum of contributions from survival and recruitment (cf. Pollard 1966).

The resulting stochastic kinship model, for kin of type **k**, is

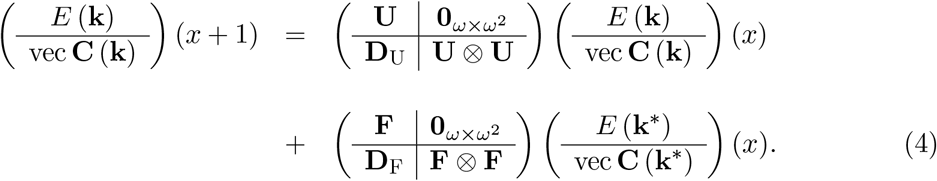

Of course, the fertility term is replaced by vector of zeros for kin types with no recruitment. Equation (4) can be written in the reassuringly familiar form

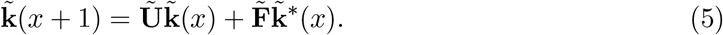

The matrices **U** and **F** in the upper left quadrant of Ũ and 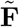 operate on the expected age distributions at age *x* to produce the expected distributions at *x* + 1, just as in equation (1).

The matrices **U** and **F** in the upper left corner of the matrices project the expected kinship vector, exactly as in the deterministic model in equation (1). The Kronecker products **U** ⊗ **U** and **F** ⊗ **F** project the covariance of the kin vector under the linear transformations given by **U** and **F**; see Appendix A.1.^3^

The matrices **D**_U_ and **D**_F_ in the lower left corner of the matrices account for the contribution to the covariance of the demographic stochasticity in transitions/survival and reproduction, respectively. Consider an individual in stage *j* at age *x* of Focal. At *x* + 1, this individual will have moved to a different stage, or died. The distribution of fates is multinomial, and the covariance matrix of the multinomial distribution is

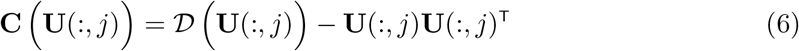

(Caswell, 2001, Eq. 15.101). In the age-classified case, there is only a single positive probability in the multinomial. The formula applies equally to stage-classified life histories (Caswell, 2001).

By age *x* + 1, the individual in stage *j* of kin **k**^*^ will have produced offspring according to *F*. Assuming that all offspring enter age class 1 and that fertility follows a Bernoulli distribution (i.e., one offspring at most), then

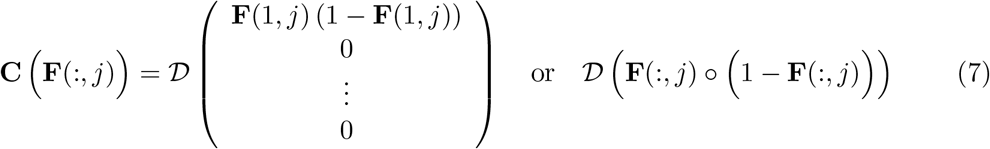

In terms of these matrices,

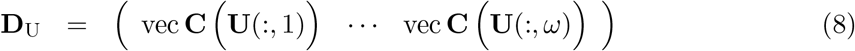

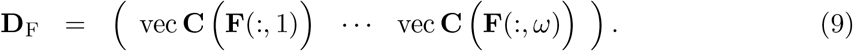

Equation (4) is essentially Pollard’s model (Pollard, 1966, eqn. 26) extended to include immigration (similar to his equation (38)), but with the immigration accounted for by the reproduction of the linked population **k**^*^(*x*). It requires only the condition that recruitment from kin of type **k**^*^ be independent of the transitions of kin of type **k**. This already follows from the independence of individuals in a branching process.

### 2.4 Initial conditions for the stochastic model

Initial conditions for the stochastic model, which include both means and covariances of kin at the birth of Focal, fall in four categories.

1. We know that Focal is alive at her birth, so the initial condition ***ϕ***_0_ for Focal contains exactly one individual in the first age class,

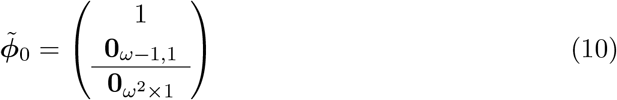
2. Focal has exactly one mother at her birth. The mother’s age is distributed according to π. A mother selected from π has a multinomial distribution, with

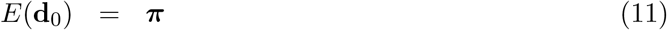

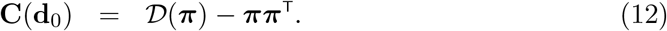

Thus

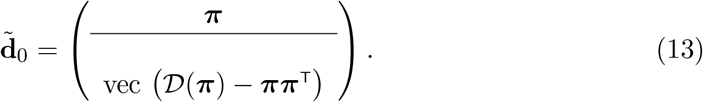
3. For those kin for which the initial condition is fixed at zero (daughters, granddaughters, great-granddaughters, younger sisters, nieces via younger sisters) both the mean and the covariance vector are zero. See Table 1)
4. Initial conditions for the remaining kin types (grandmothers, great-grandmothers, older sisters, nieces via older sisters, aunts older than mother, aunts younger than mother, cousins from aunts younger than mother, and cousins from aunts older than mother) are mixtures, over the distribution of age at maternity, of some other type of kin.

For example, the older sisters at Focal’s birth are the children of Focal’s mother at the birth of Focal. As in the deterministic model, the initial condition for the mean of older sisters is the mean of children over the mixture distribution of mothers ages.

Let π be the distribution of age at maternity. If the initial condition for kin of type **k** (e.g., older sisters) is a mixture of kin of type **k**^*^ (e.g., children) then

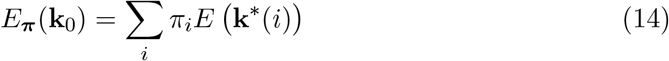

The covariance of the initial population is calculated from the ‘law of total covariance’ for the covariance of a mixture (Appendix A.2):

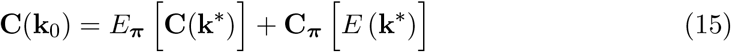

where

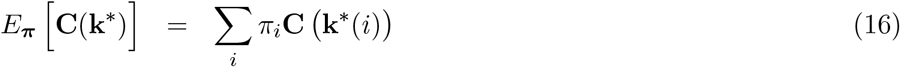

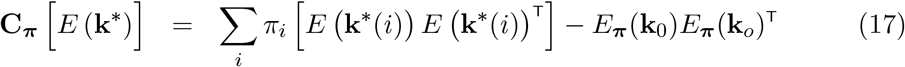

For ease of reference, we define the initial condition for kin of type **k** as a function ℐ (*·, ·*) of the mixing distribution and the source kin type **k**^*^,

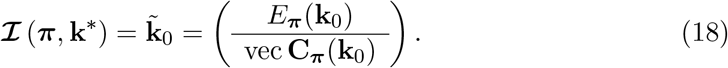

### 2.5 Summary of the model

Given the new definitions of the kinship vector, the survival and fertility matrices, and the initial conditions, the stochastic model follows closely the structure of the deterministic model. Table 2 summarizes the calculations for the stochastic model. For comparison, Table 1 shows the structure of the corresponding deterministic model (Caswell, 2019).

**Table 2:**
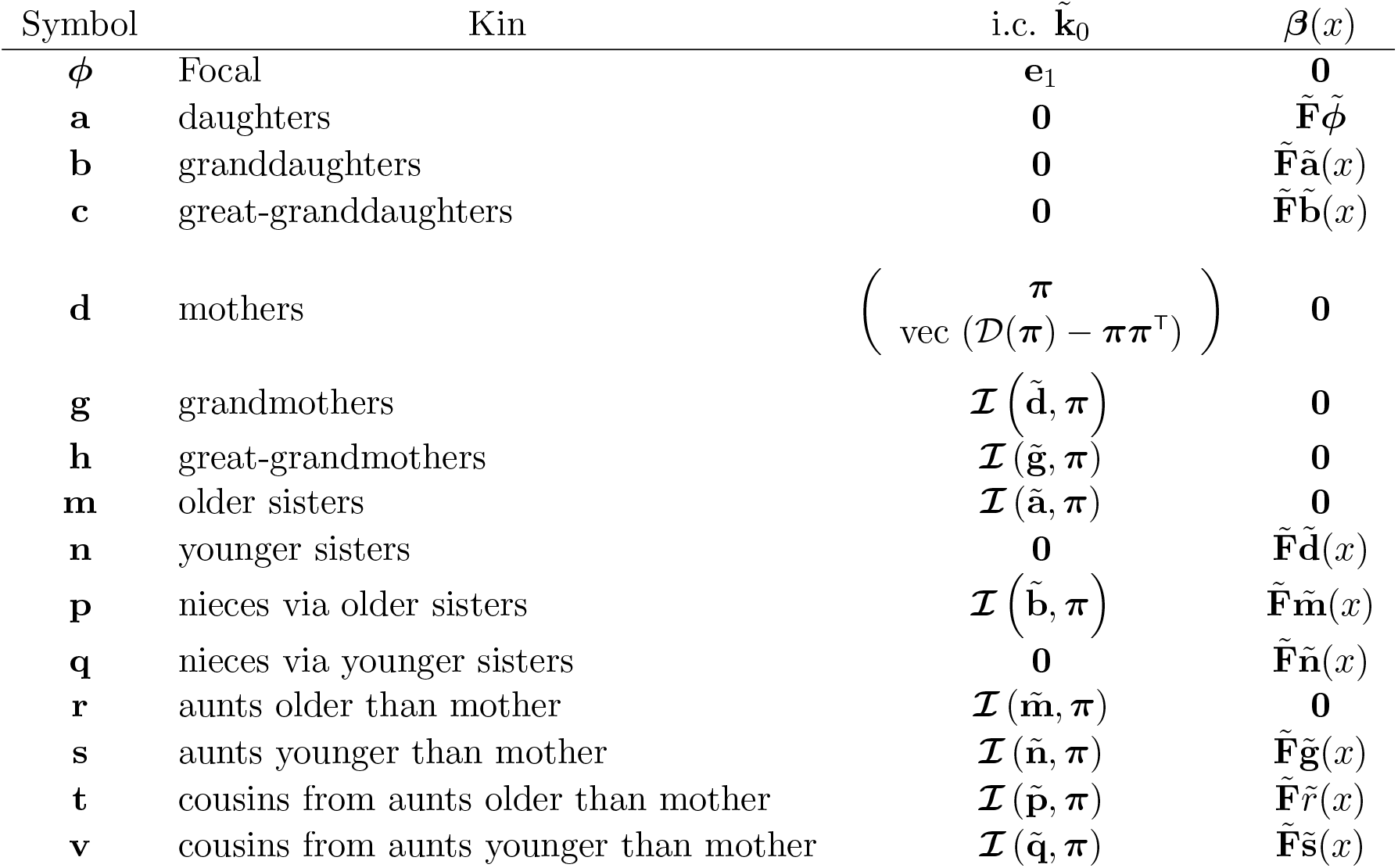
The initial condition and the recruitment term for each type of kin in the stochastic model (one-sex, time-invariant). The initial condition function ℐ (*·, ·*) is given by equation (18). The initial condition for mothers is given in (12).

## 3 Outputs of the stochastic kinship model

In addition to the vector 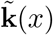 it is important to consider other outputs that are functions of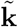. Examples include linear transformations of the kin vector, such as numbers or weighted numbers of kin, numbers of kin with some kind of condition, and approximate two-sex results.

They require some care in accounting for stochasticity.

### 3.1 Linear transformations of the kin vector

Consider an output **y** that is a linear transformation of a kin vector **k**, written as

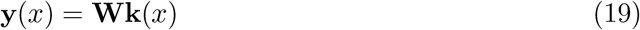

for some matrix **W**. The expected value and the covariance matrix of the output **y** are

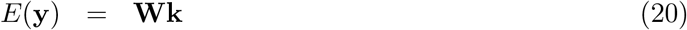

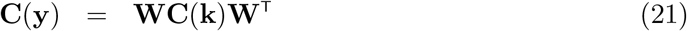

so that

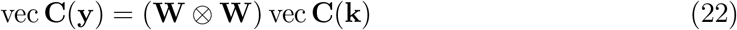

The block-structured output vector is

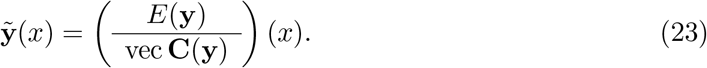

The combined dynamics of both the kin vector and the output vector are given by^4^

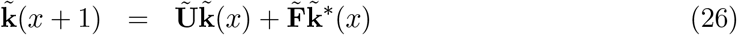

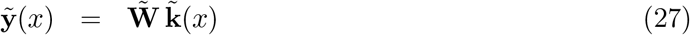

where

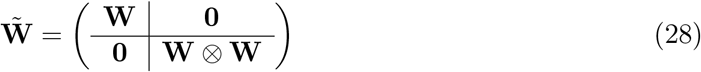

Equations (47) and (48) can be called the state equation and output equation, respectively.

Some examples of linear transformations:

- Total numbers of kin. In this case **y** is a scalar, and the transformation matrix is

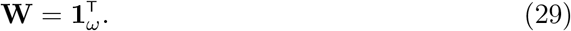

and

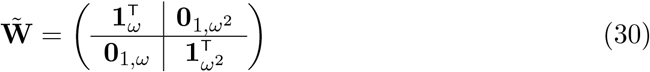

The expected number is the sum of the expectations of each age class, and the variance in total number is the sum of all the variances in, and covariances among, the age classes.
- Weighted numbers of kin. The vector **1** can be replaced with any vector **c**, weighting the kin of age class *i* with the value *c*_*i*_ (e.g., per-capita income). The output matrix is

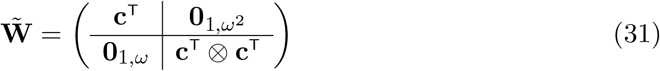

The mean weighted number is the weighted sum of the mean numbers. The weighted covariance structure is produced by the vector **c**^T^ ⊗ **c**^T^, which provides the proper weights for each variance and covariance.
- Multiple selected ages of kin. Suppose we want the numbers of kin in several different age ranges. Dependency ratios, for example, are calculated from the numbers of individuals in young dependent, independent, and old dependent age ranges.

For example, suppose that of six age classes, classes 1 and 2 are young dependent, 3 and 4 are independent, and 5 and 6 being old dependent. The vector **y** is 3 × 1 and the transformation matrix is

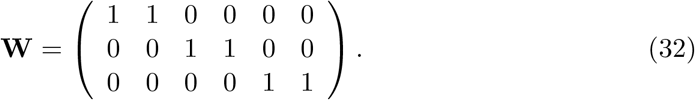

and the output matrix is given by equation (28).

### 3.2 The GKP factors: independent realizations of kin

The one-sex matrix kinship model projects female kin through female lines of descent (Caswell, 2019). In the absence of a complete set of two-sex rates, a useful approximation is obtained by multiplying the mean kin vector by a factor, introduced by Goodman, Keyfitz, and Pullum (1974) and called the ‘GKP factors’ by Caswell (2022), specific to each type of kin. The factors are based on an androgynous approximation in which males and female rates are identical (see Caswell 2022 for more discussion). The GKP factors for each type of kin are given in Table 3.

**Table 3:** The GKP factors used to transform female kin through female lines of descent to both sexes through all lines of descent. Reproduced from Caswell, Verdery, and Margolis (2023).

| Transformation | GKP factor |
| --- | --- |
| daughters $\mapsto$ children | 2 |
| granddaughters $\mapsto$ grandchildren | 4 |
| great granddaughters $\mapsto$ great grandchildren | 8 |
| mothers $\mapsto$ parents | 2 |
| grandmothers $\mapsto$ grandparents | 4 |
| great grandmothers $\mapsto$ great grandparents | 8 |
| sisters $\mapsto$ siblings | 2 |
| nieces $\mapsto$ nieces-nephews | 4 |
| aunts $\mapsto$ aunts-uncles | 8 |
| female cousins $\mapsto$ cousins | 8 |

The covariance structure of the GKP-weighted kin describes the result of independent realizations of a number, given by the GKP factor, of independent lineages. Grandchildren, for example, arise from four lineages (daughters of daughters, daughters of sons, sons of daughters, and sons of sons). Given the independence of individuals, the covariance of the sum of these four lineages is

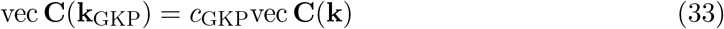

where *c*_GKP_ is the appropriate GKP factor.

## 4 An example of the stochastic model: Japan over seven decades

As an example of the stochastic model, I present some results based on demographic rates from Japan in 1947 and in 2019. Japan underwent dramatic demographic changes over this period. In 1947, mortality was high (period life expectancy at birth of only 53.7 years) and fertility was also high (period TFR of 4.6 children per woman). By 2019 both mortality and fertility had declined: life expectancy had increased to 87.4 years and TFR had shrunk to only 1.3 children per woman. These changes have opposing effects on the kinship network. Low fertility means that fewer kin are produced by birth, but low mortality means that fewer kin are lost by death. Some of the consequences for kin were explored by Caswell (2019). Here we use these contrasting situations to examine specifically stochastic elements of the kinship network.

This analysis is intended as an example, and not a serious exploration of kinship networks in Japan. The analysis is based on period mortality rates from the Human Mortality Database (HMD, 2022) and period fertility rates from the Human Fertility Database (HFD, 2022).

**Figures displaying results:** This analysis creates a large number of graphs. For convenience of the reader, these are collected in a Section 7 at the end of the paper. This makes it possible to compare across kin types, ages of Focal, and years, and to explore some ways of displaying results. The reader will no doubt think of more.

### 4.1 Kin numbers: Means, variances, and standard deviations

Figure 3 presents the mean, variance, and standard deviation of GKP-weighted numbers of kin, of each of the types in Figure 1 (with older and younger kin, for example older and younger sisters, combined).

**Figure 3:**
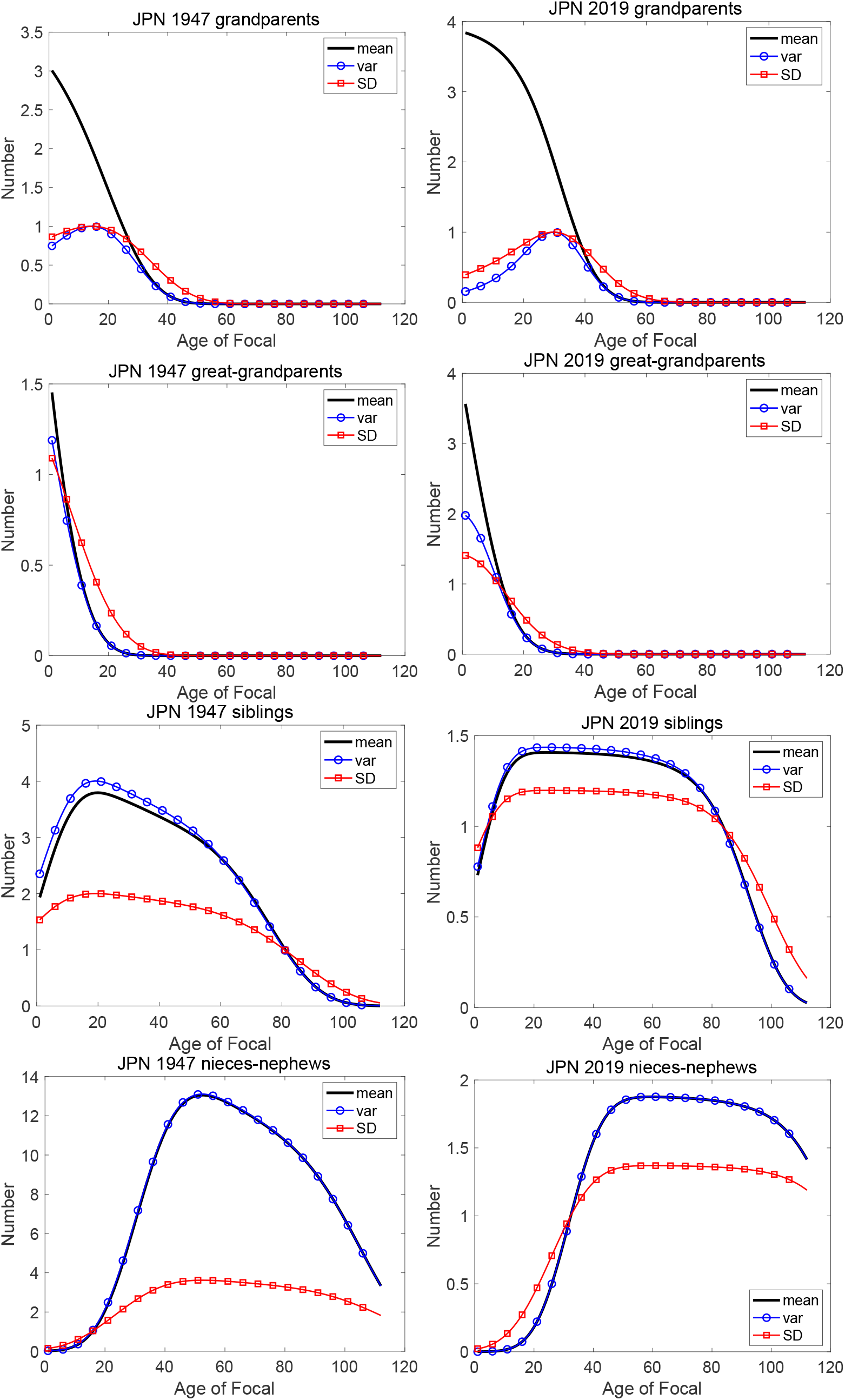
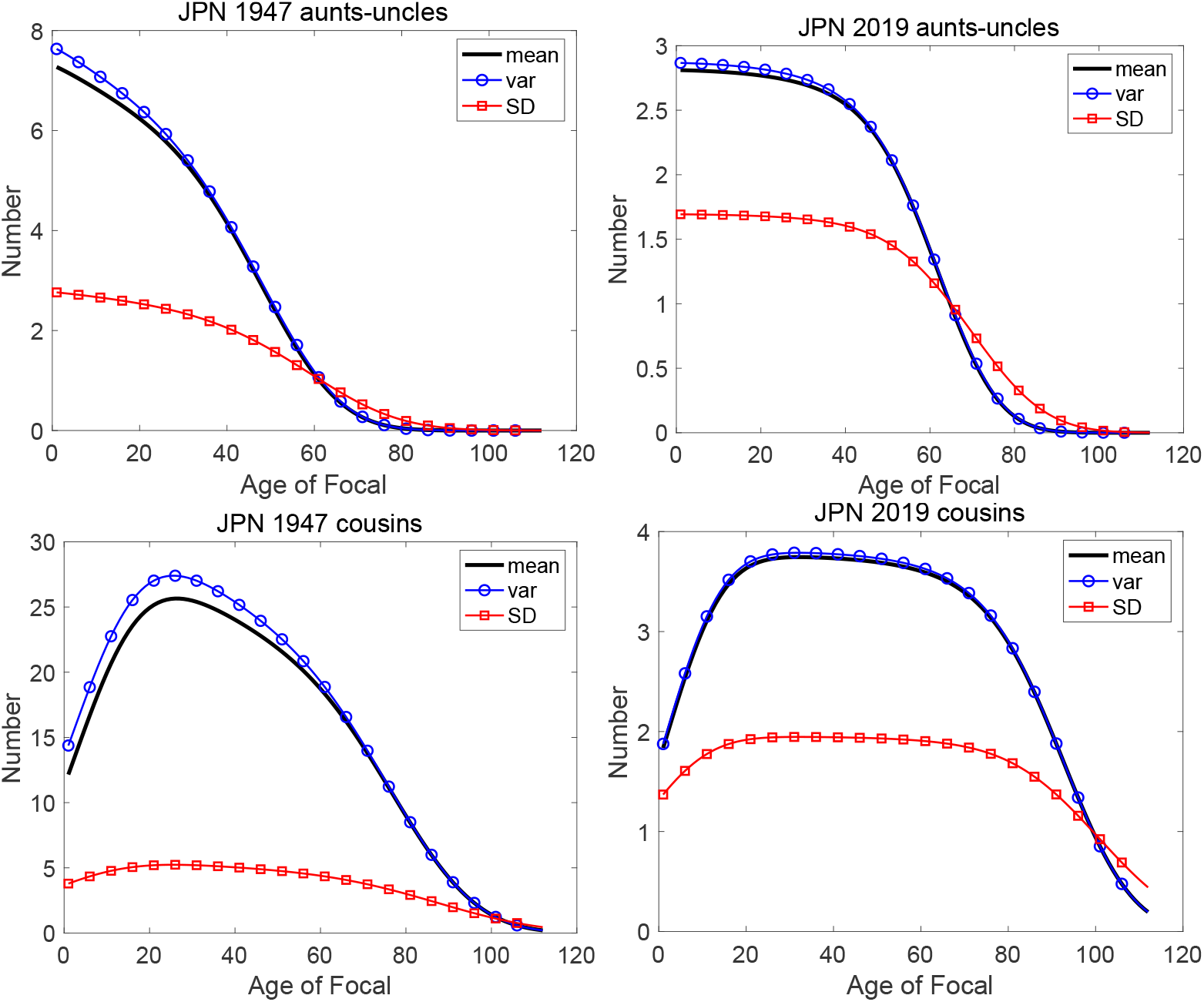
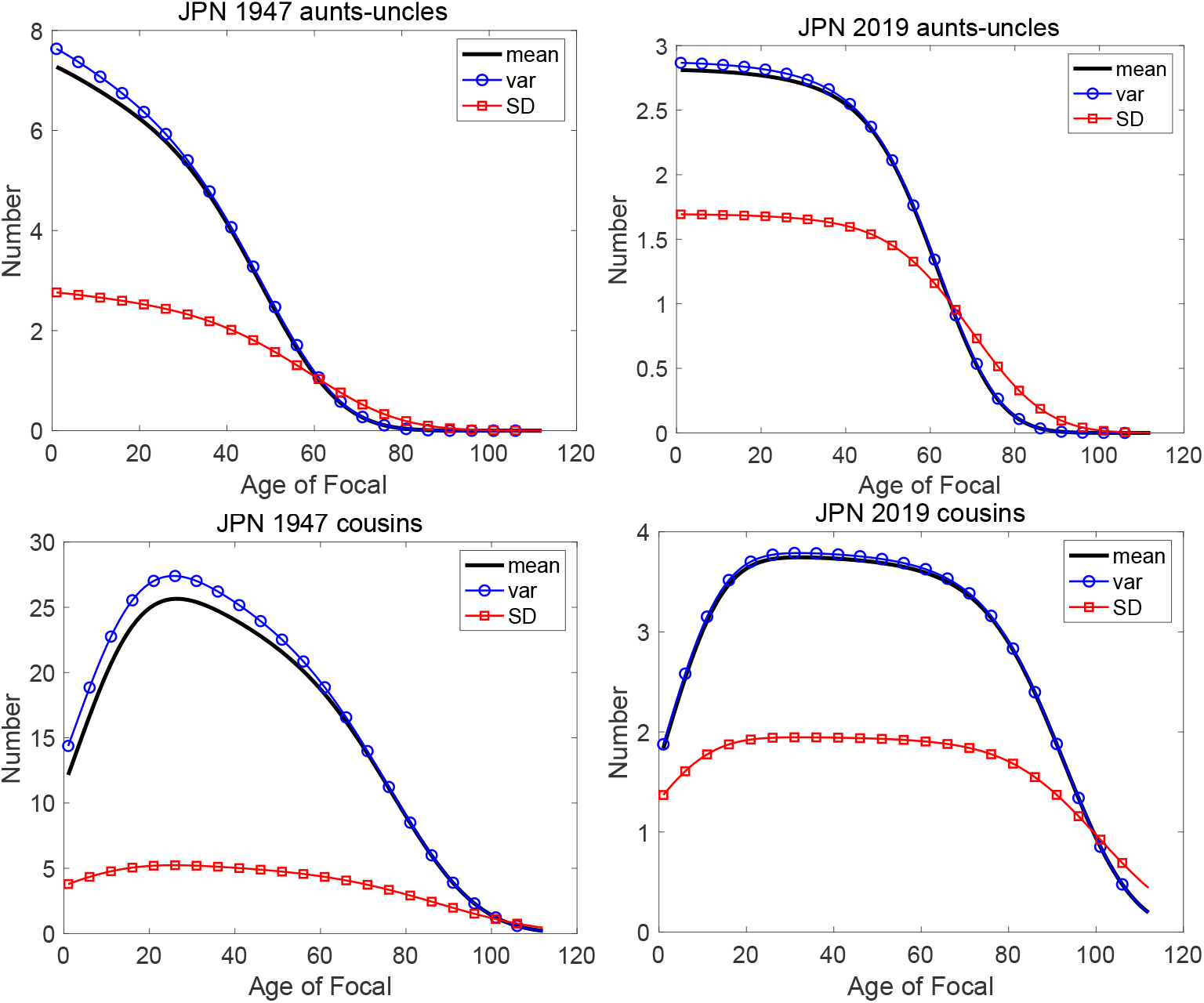
(Part 1) The mean, variance, and standard deviation (SD) of the number of kin, of each type, as a function of the age of Focal, under the rates for Japan in 1947 and 2019. Note different scales on the y-axis.

- The patterns of the mean kin numbers over the ages of Focal are familiar from the comparisons of Japanese kinship in Caswell (2019).
- On average children, grandchildren, and great-grandchildren are more numerous in 1947 than in 2019 Under 1947 rates, Focal would see a rapidly growing family of descendants, whereas under 2019 rates subsequent generations would decline.
- Parents, grandparents, and great-grandparents die off more rapidly under 1947 rates, and Focal has on average fewer grandparents and great-grandparents at birth under 1947 rates.
- Siblings, nieces-nephews, aunts-uncles, and cousins are more abundant (sometimes much more abundant; see cousins) under 1947 rates than under 2019 rates.
- The mean numbers of kin are accompanied by considerable variation, as expected. For example, under 1947 rates Focal averages about 4 siblings at age 20, with a standard deviation of about 2. Under 2019 rates Focal averages about 1.4 siblings, with a standard deviation of about 1.2
- More interesting and more useful is the variance in kin numbers. For all kin other than direct ancestors (parents, grandparents, great-grandparents), the variance is close to the mean (sometimes indistinguishable from the mean in these plots; e.g., for nieces-nephews). This is compatible with a Poisson distribution for kin numbers, and suggests the Poisson as a good approximation to the distribution of the numbers of these kin (Stuart and Ord, 1987, Sec. 3.34).
- The variance in numbers of ancestors shows a different pattern. The variances in the numbers of parents, grandparents, and great-grandparents are distinctly underdispersed relative to the mean. At birth, Focal has exactly 2 parents, so the variance at age *x* = 0 is zero. Focal has at most 4 grandparents, and 8 great-grandparents. The living ancestors at subsequent ages of Focal are the survivors of those at Focal’s birth. Thus a binomial distribution is an appropriate description for the these kin. see Section 4.2.

### 4.2 Prediction intervals and uncertainty

From the mean and variance a prediction interval for the number of kin can be calculated; this interval captures a specified percentage of the variation among Focal individuals subject to the rates under study.^5^ This requires a distribution; we can estimate that distribution from the means and variances. The distribution of kin numbers is discrete, with support on the non-negative integers. Those kin for which the variance is close to the mean are compatible with the Poisson distribution. This distribution has often been assumed for kin numbers even without knowledge of variance (e.g., Song, Campbell, and Lee, 2015; Song and Mare, 2019; Caswell, Verdery, and Margolis, 2023; Feng, Song, and Caswell, 2023).

From Figure 3 we know that the variance in kin number is close, sometimes very close, to the mean for all kin types except parents, grandparents, and great-grandparents. Those kin are the survivors of some initial number at the birth of Focal, with no subsequent recruitment. Thus the binomial distribution is an appropriate for these. The initial number is 2 for parents, 4 for grandparents, and 8 for great-grandparents.^6^ The prediction interval for a discrete random variable, for some probability *p*, is obtained by finding the smallest integer kin number such that the cumulative distribution function (Poisson or binomial) evaluated at that kin number is greater or equal to *p*, using the Matlab functions `poissinv` or `binoinv`.

Figure 4 shows the 90% prediction interval (i.e., the interval between the 5th and the 95th percentiles of the distribution of kin numbers) and the 50% prediction interval (i.e., the interval between the 25th and 75th percentile, also known as the inter-quartile range (IQR)).

**Figure 4:**
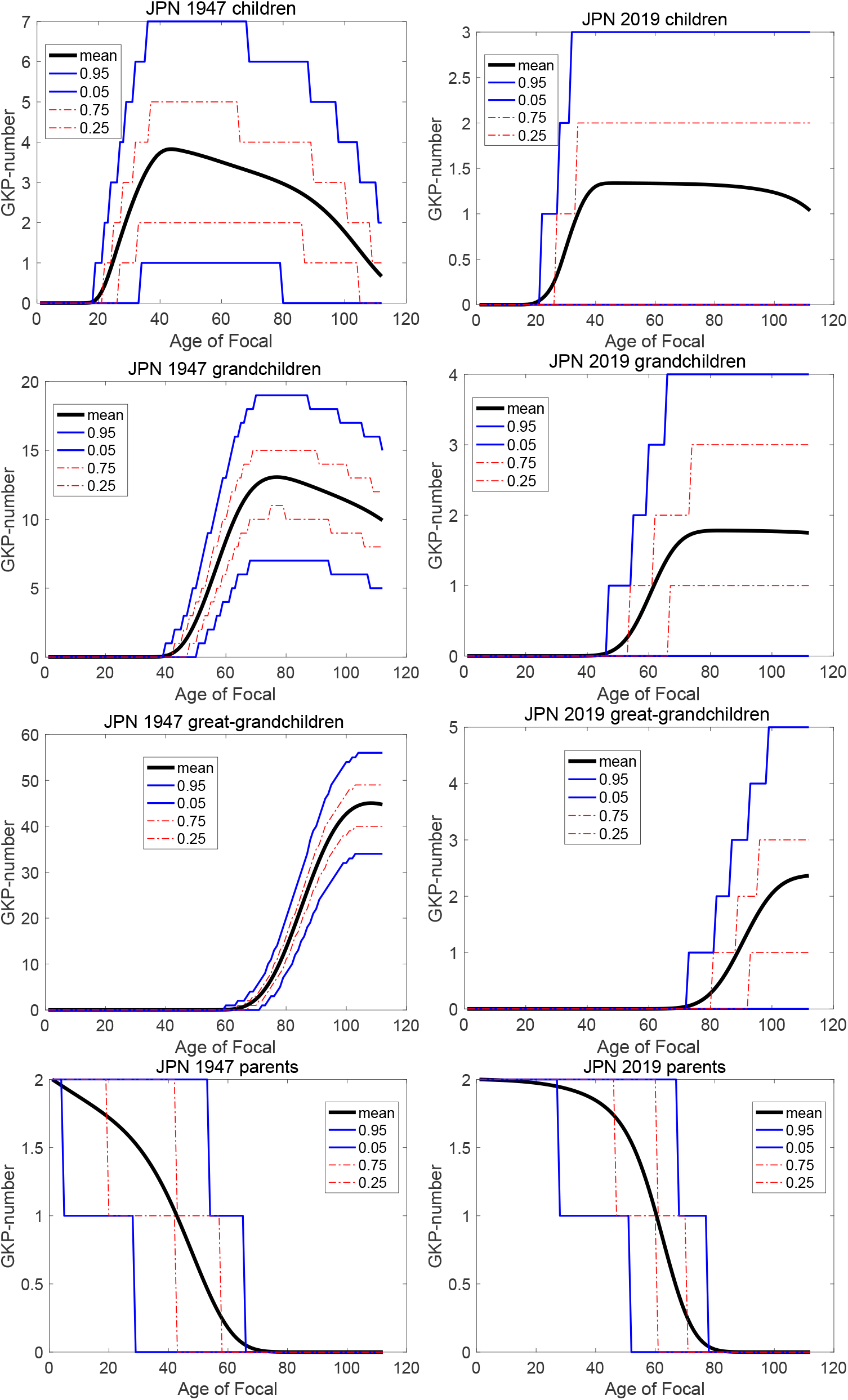
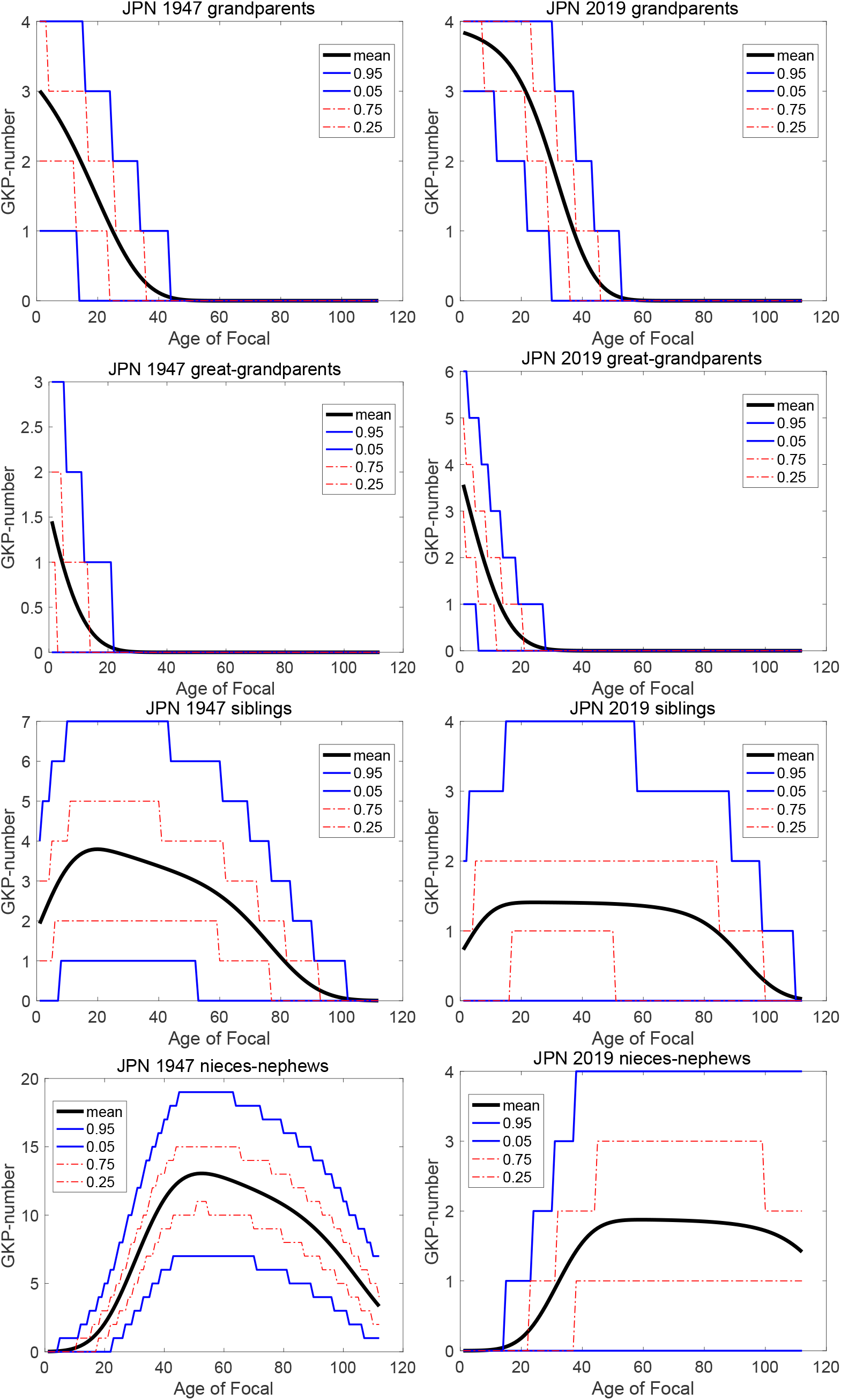
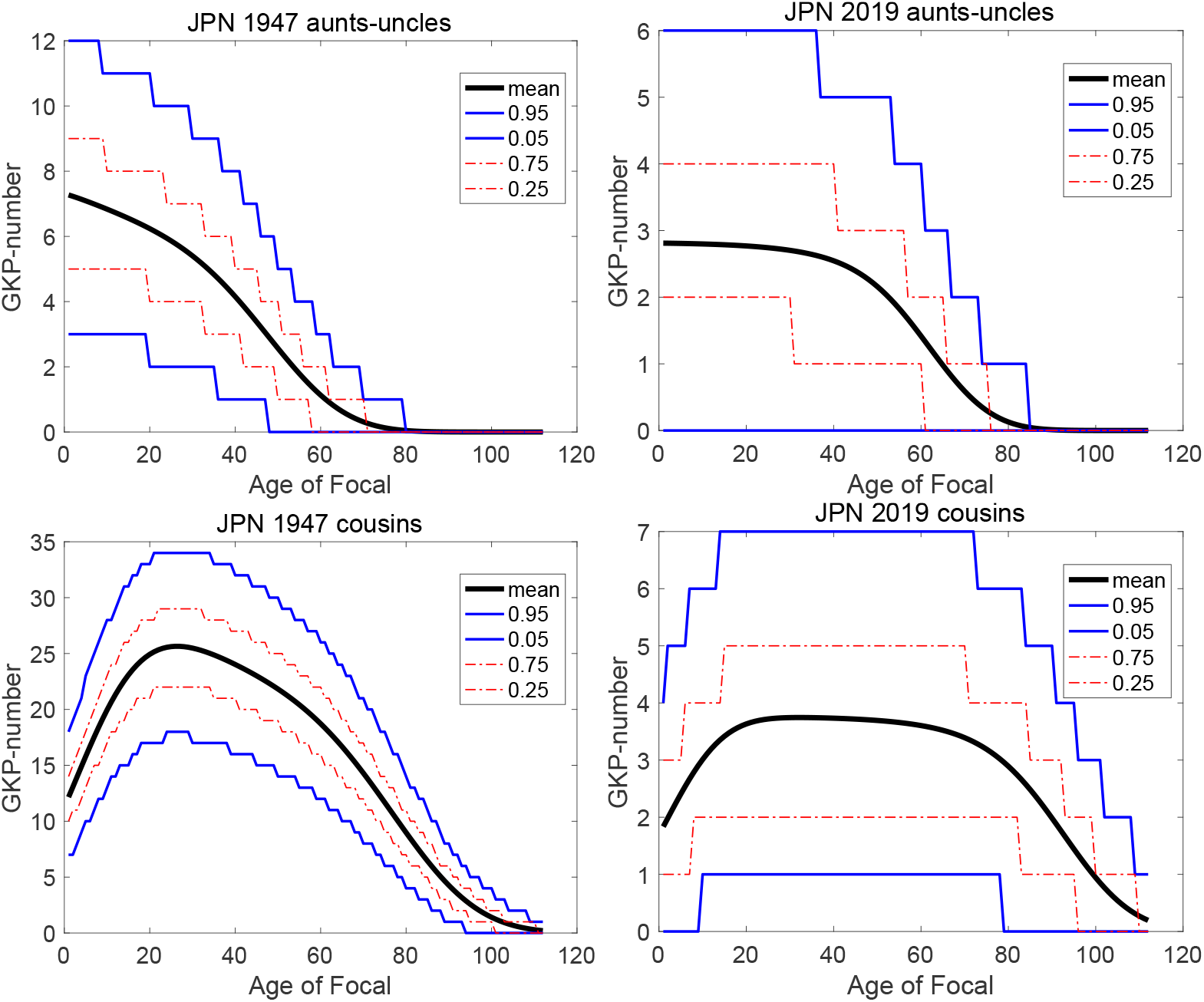
(part 1) Expected number of kin, of each type, with 5%, 25%, 75%, and 95% prediction intervals.

- The numbers of children, grandchildren, and great-grandchildren have wide prediction intervals. Under 2019 rates, Focal at age 60 would expect to have about 1.5 children at age 60. Ninety percent of Focal individuals subject to these rates would have between 0 and 3 children; 50% would have between 0 and 2 children. Under 1947 rates the expected number of children at age 60 is about 3, with a 90% prediction interval from 1 to 7.
- Some of the differences between years in the prediction intervals are impressive. For example, under 1947 rates, at end of her life Focal could expect about 45 great-grandchildren, with an IQR of [40, 50]. Under 2019 rates, Focal could expect only 2.5 great-grandchildren, with an IQR of [1, 3].
- Knowledge of the prediction intervals, and hence of the variation to be expected among individuals in the numbers of their kin, may be useful for planning of family services. This also opens the potential for comparison with empirical studies of kinship networks from register data (e.g., Kolk et al., 2023).

## 5 Some additional outputs

“But wait, there’s more.”

Ron Popeil, celebrity TV salesman

Some calculations that are routine under the deterministic model for expected kin require more careful consideration, and provide extra information, in the stochastic model. Three of these are considered here: calculations of presence or absence of kin, the prevalence of conditions among kin, and dependency ratios among kin.

### 5.1 Experience of the presence of kin

The probability of having at least one kin of a specified type is the complement of the zero term of the distribution of kin numbers. The probability of having at least one, say, grand-child can be interpreted as the fraction of the population that has at least one grandchild; this is sometimes a more intuitive result than the mean number of kin.

For the Poisson-distributed kin with mean kin number *k* the probability of at least one kin is

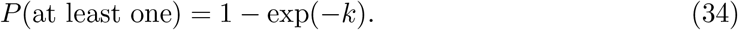

For the binomially-distributed kin (parents, grandparents, great-grandparents) with initial number *N*, the probability is

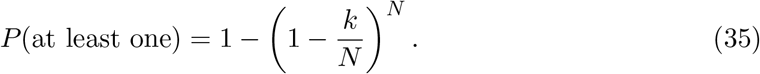

These calculations generalize easily to the proportion of individuals with at least 2 kin, at least 3 kin, and so on, using the cumulative distribution function of *k*, to provide the proportion of individuals with 2 or more kin, 3 or more kin, etc.

The probabilities of having at least one kin, and of at least two kin, are shown in Figure 5 for all kin types; the differences between 1947 and 2019 rates are not surprising given the differences in the expected number of kin.

**Figure 5:**
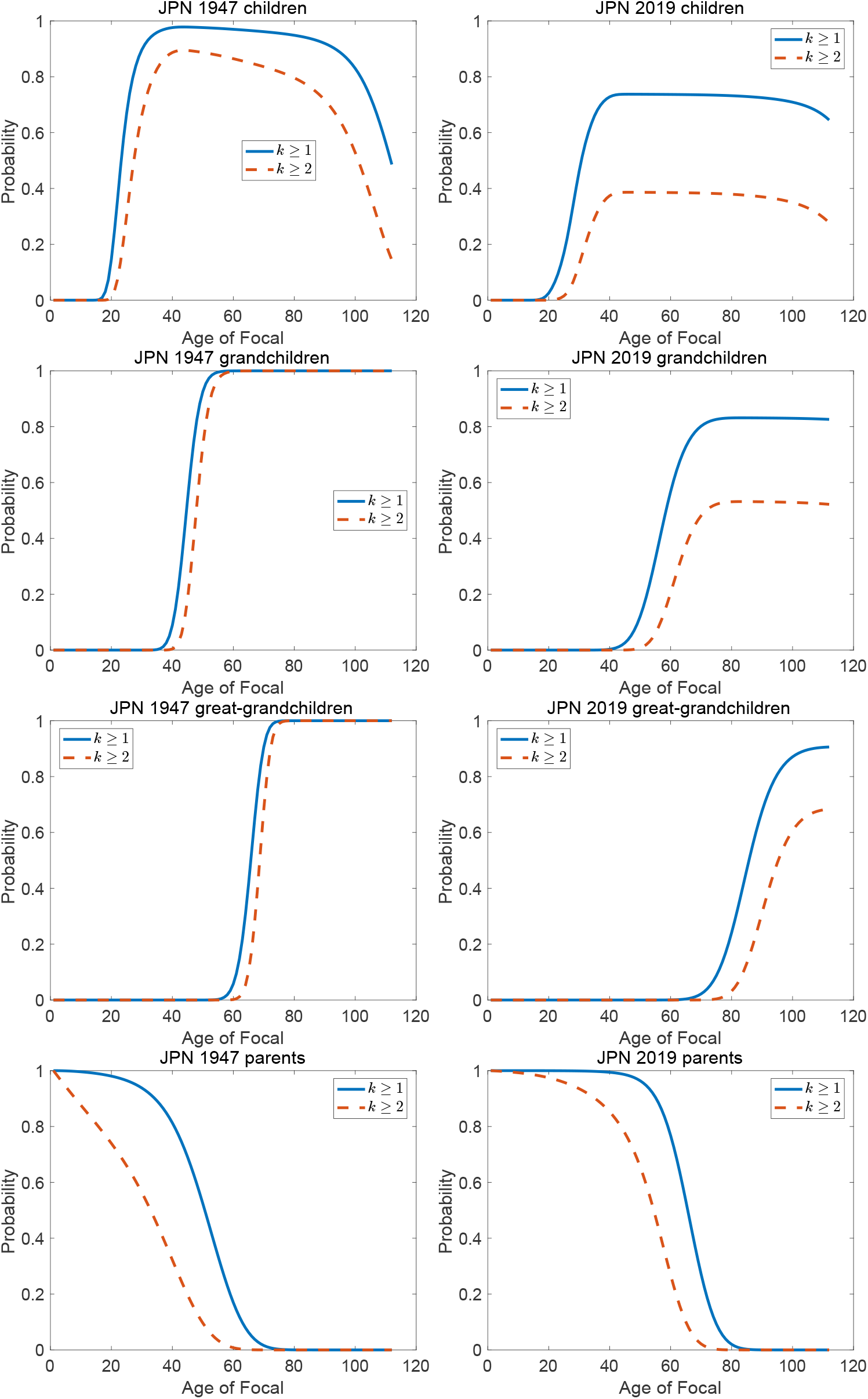
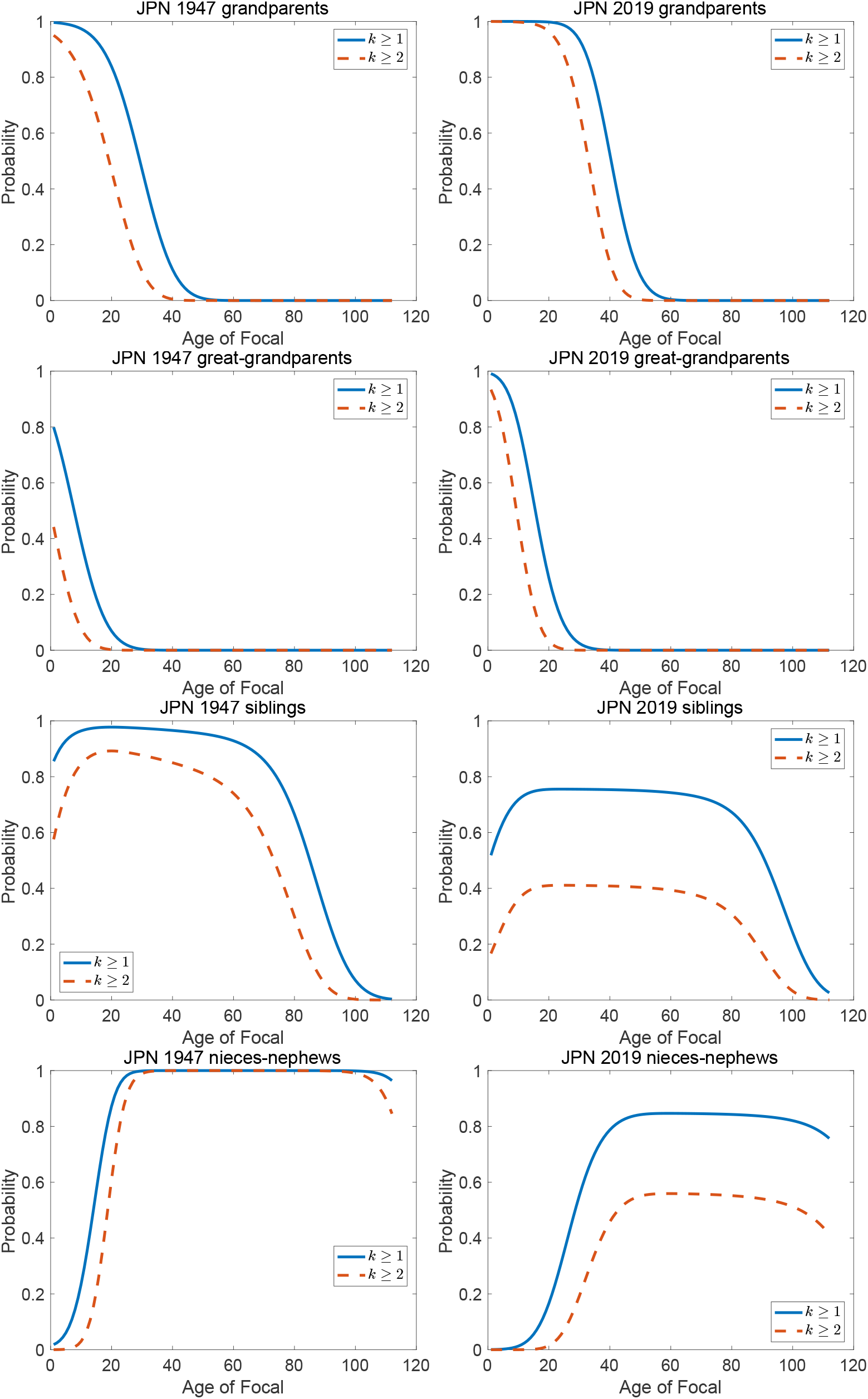
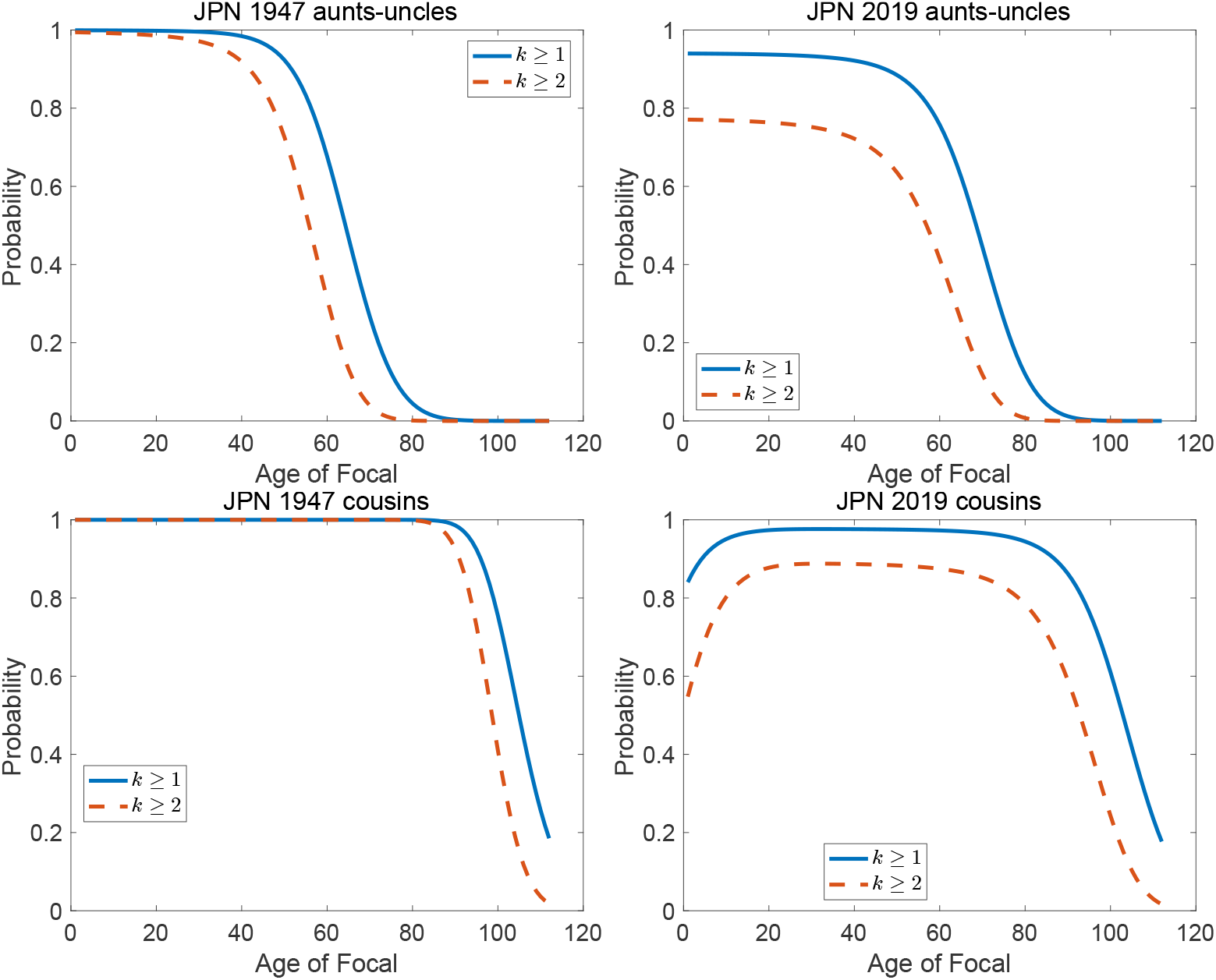
(part 1) The probabilities of having at least one, and at least two kin of each kind, as a function of the age of Focal. Probabilities calculated from binomial distributions for parents, grandparents, and great-grandparents and Poisson distributions for all others, fitted by the method of moments.

- Children and descendants. The difference between increasing and decreasing kin networks appears in the probabilities of having at least one kin.
  – Under 1947 rates, Focal at age 40 is almost certain to have at least one child, but Focal under 2019 rates has a probability of only about 0.7 of having at least one child.
  – At age60, Focal under 1947 rates is almost certain to have at least one grandchild, but Focal under 2019 rates has a probability of only 0.4.
  – At age 80, Focal under 1947 rates is almost certain to have at least one great-grandchild, but Focal under 2019 rates has only about a 20% chance.
- In contrast, Focal under 2019 rates has a higher chance of having at least one surviving parent, grandparent, and great-grandparent than does Focal under 1947 rates.
- The between-year differences are smaller for those kin that have non-zero initial populations and non-zero recruitment during the life of Focal (siblings, nieces-nephews, aunts-uncles, and cousins). The higher survival but lower fertility under 2019 rates work in opposite directions.
- The pattern for cousins is interesting. On average, Focal in 1947 has up to 25 cousins, whereas Focal in 2019 never averages more than four. This translates into differences in the probability of at least one cousin: Focal in 1947 is almost certain to have at least one cousin at every age from birth up to age 90. Focal in 2019 has a high probability, but not certainty, of having at lest one cousin up to age 60, at which point the chance begins to decline rapidly.
- Differences in mean numbers of kin need not be directly proportional to differences in the probability of at least one kin. If the mean number is very small, however, it is nearly equal to the probability of having at least one (because, for *x* small, 1 − exp(−*x*) ≈ *x*).
- The difference between the probability of at least one and at least two kin is amplified under 2019 rates, because the lower fertility in 2019 reduces the recruitment of new kin. For example, under 1947 rates, Focal is almost certain to have at least two grandchildren after the age of 50. But under 2019 rates, the probability of at least one grandchild only reaches about 0.8, and the probability of at least two only reaches 0.5. Similarly dramatic patterns appear for nieces-nephews.

### 5.2 Prevalence and incidence: proportions or probabilities?

The prevalence of a condition is the fraction of a population exhibiting that condition. Sometimes the condition is that of receiving a diagnosis of a disease; the fraction of the population experiencing this event is called, confusingly, the incidence rate. Below, we will see an analysis of cancer incidence in Japan. Prevalences have no dynamic content; they are measured at a single point in time, do not include longitudinal data following individuals, and hence are blind to individual histories (Caswell and Zarulli, 2018).

#### 5.2.1 Prevalence of multiple conditions

Age-specific prevalence schedules are often standardized against an age structure, giving a scalar value of age-standardized prevalence: the mean number of individuals exhibiting the condition. Age-standardized prevalences or incidences can be standardized against any age distribution, in particular the age distributions of any of the types of kin of Focal. Mean age-standardized number of kin have been calculated for unemployment by Song and Caswell (2022) and for dementia by Feng, Song, and Caswell (2023, 2024).

To calculate covariances we recognize that age-standardized prevalences are linear transformations of the kin vector. Let **W** be a matrix of dimension *c* × *ω*. Each of the *c* rows contains the age-specific prevalence schedule of one of *c* conditions. As prevalences, it makes sense to assume that the column sums of **W** are less than or equal to 1.

For a single binary condition, *c* = 1, and the linear transformation matrix **W** contains a single row. We will use this in an example where the condition is a new cancer diagnosis.

For multiple alternative conditions, *c* > 1 and **W** is *c* × *ω*. For example, NCD Risk Factor Collaboration (2024) published a worldwide analysis of trends in body weight, reporting the adult prevalence of underweight and of obesity based on BMI data. As these conditions are defined, an individual can fall in only one of the categories. To analyze these conditions, *c* might equal 3, and the first row of **W** could contain the age-specific prevalence of underweight, the second row the prevalence of obesity, and the third row the prevalence of neither.

Multiple conditions, with *c* > 1, can also result from the joint occurrence of combinations of independent binary factors. For example, Palmqvist et al. (2023) studied the prevalence of three measurable conditions associated with dementia.^7^ They report age-specific prevalences of all 8 combinations of presence and absence of the three conditions. These results could be used to calculate the means, variances, and covariances of the number of individuals exhibiting each of the eight combination of conditions.

#### 5.2.2 Proportions or probabilities?

The stochastic model introduces an important new distinction: prevalence may be interpreted as either a fixed proportion or as a probability. The fixed interpretation says that exactly the proportion **W**(*i, j*) of age class *j* exhibits condition *i*. The random interpretation says that members of age class *j* either exhibit or do not exhibit the condition, with probabilities **W**(*i, j*) and (1*−***W**(*i, j*), respectively.^8^ The distinction between fixed and random prevalence makes no difference in calculating mean age-standardized numbers, but does for calculating variances and covariances. In the case of a random prevalence, the variance in the age-standardized numbers exhibiting the condition reflects both the covariance generated by the branching process for 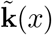 and the variance generated by the random nature of the prevalence.

The matrix **W** defines a linear transformation from the kin vector **k** to an output vector **y** giving the means and covariances of the age-standardized number of individuals with the condition. Thus, in exact analogy to the equations (4) the output equation becomes

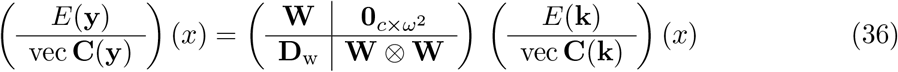

The matrix **D**_w_, just as in equation (4) captures the contributions of the stochasticity in the transition contained in **W** to the covariance of **y**.

1. For fixed prevalences, there is no stochasticity; every individual in age class *i* produces exactly *w*_*i*_ individuals with the condition. In this case,

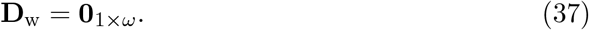
2. When prevalence is random,

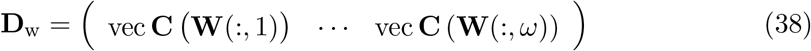

(cf. equation (8)). When there is only one condition, *c* = 1 and the distribution of the number of individuals exhibiting the condition is Bernoulli. Let *w*_*j*_ = **W**(1, *j*). Then

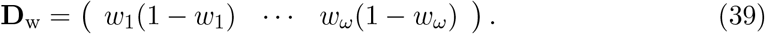

#### 5.2.3 An example: Cancer incidence in Japan

The incidence of new cases of cancer has increased in Japan in recent years (Katanoda et al., 2021). Age-specific incidence rates are not available as far back as 1947, but as an example the rates^9^ for 1975 and 2018 have been applied to the kin age distributions under 1947 and 2019 rates, respectively. Rates are available for a variety of cancers; results shown here are for all cancers combined. These rates provide the prevalence of experiencing a first cancer diagnosis at each age. Figure 2 shows the rates as a function of age.

The results of the analysis are shown in Figure 6, giving the mean number of kin, of each type, experiencing a cancer diagnosis at each age of Focal.

**Figure 6:**
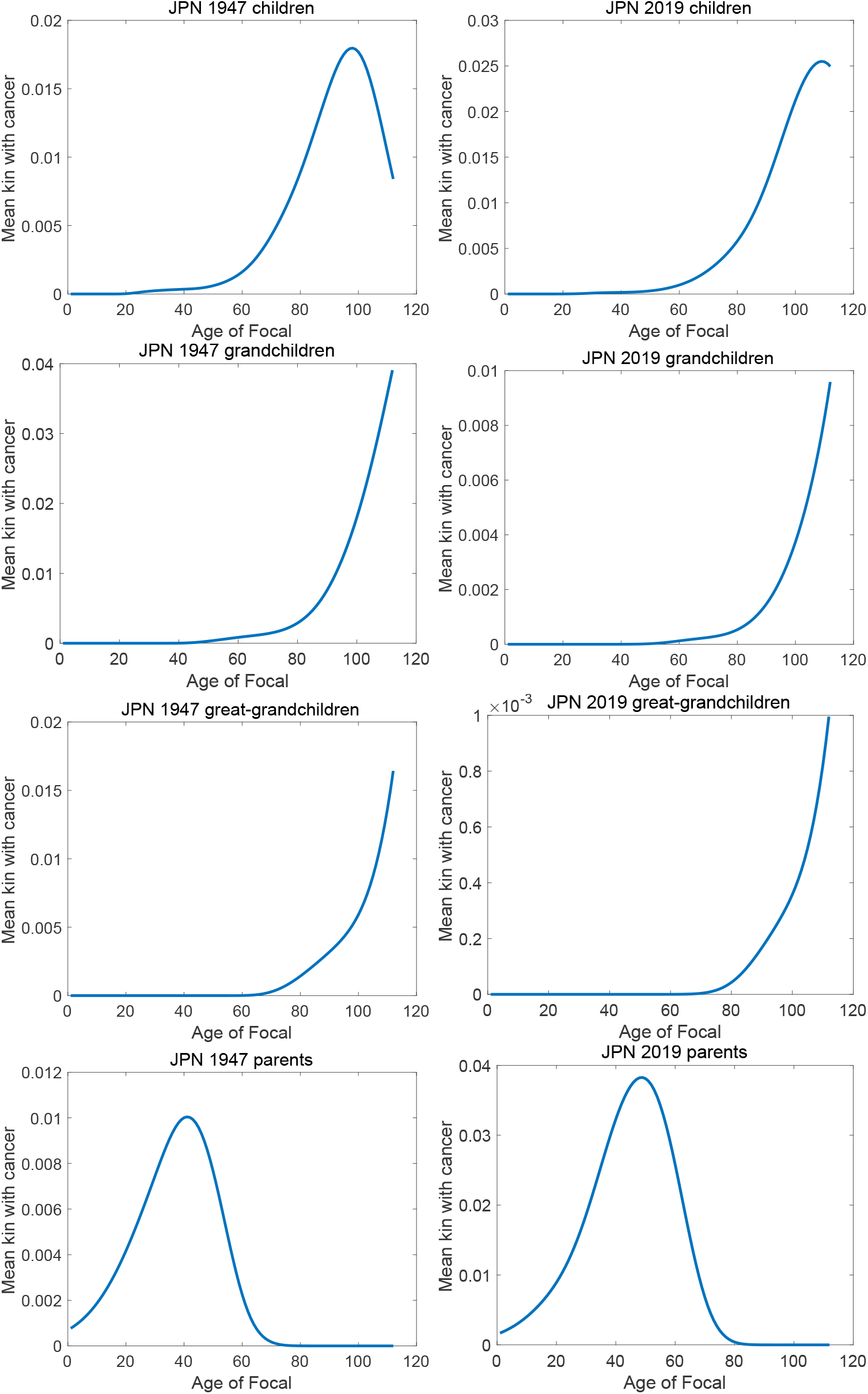
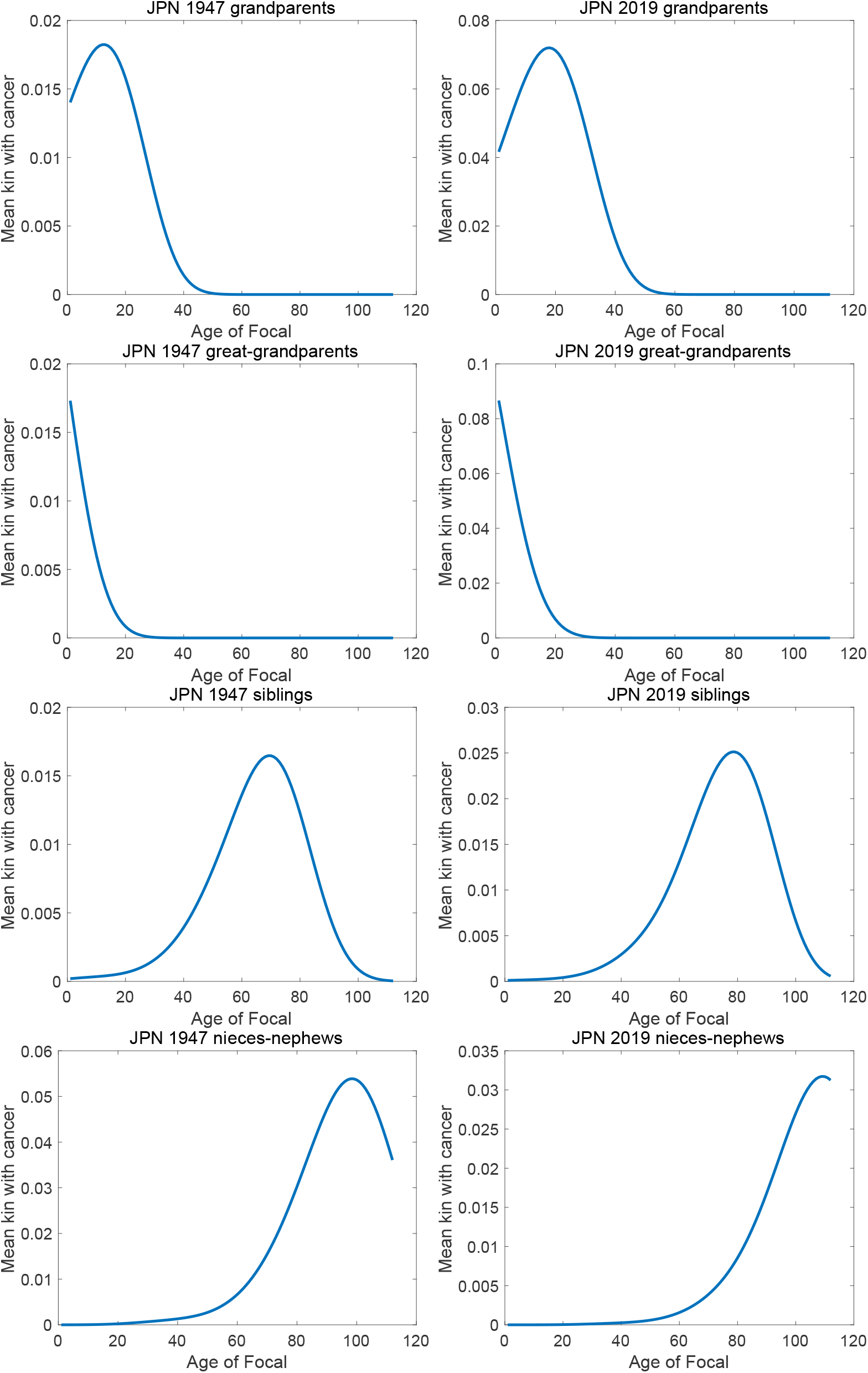
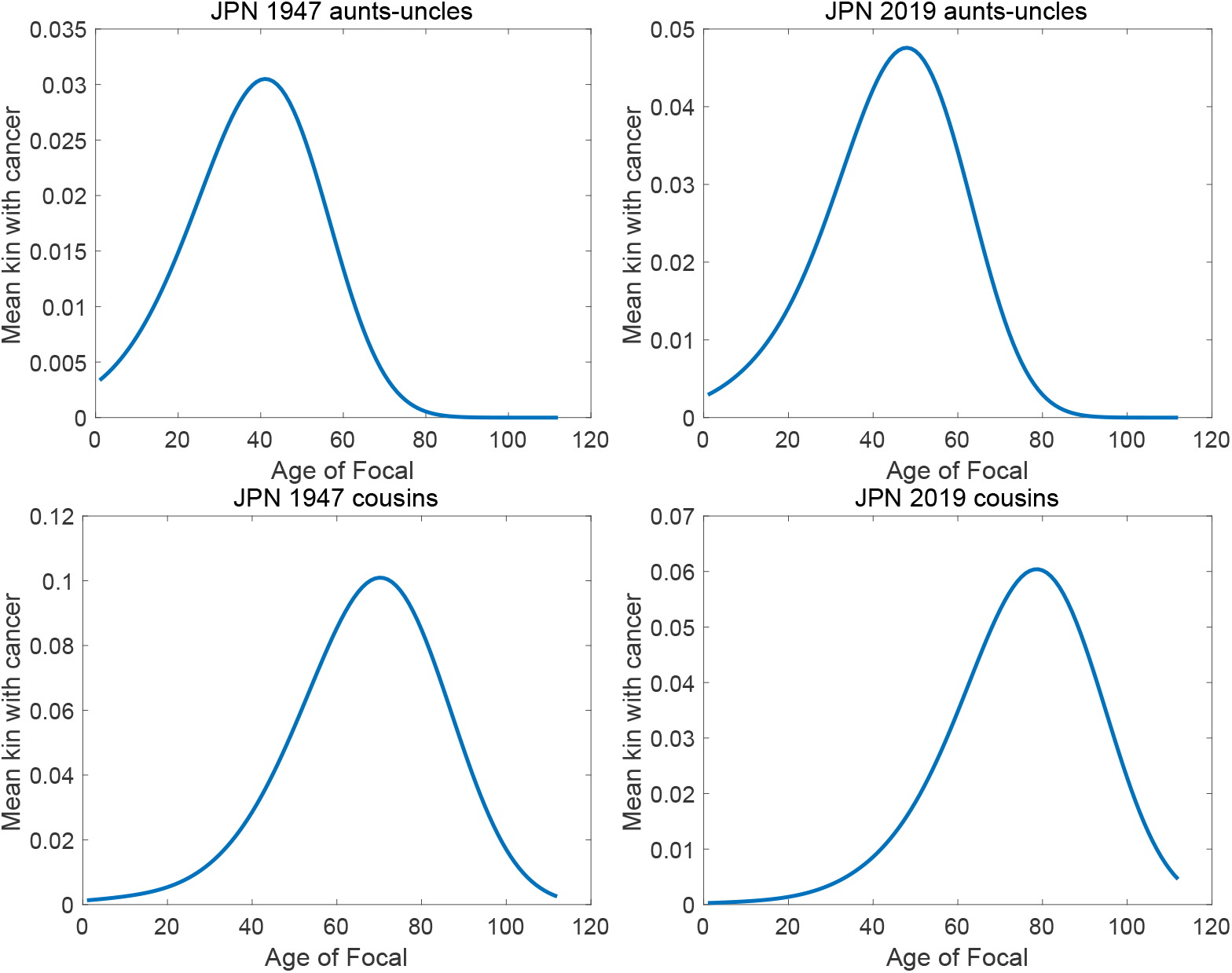
(part 1) Expected number of kin experiencing a cancer diagnosis (‘incidence’) as function of the age of Focal. Note the different scales on the y-axis.

- The variances of the age-standardized prevalence are not shown in Figure 6 because for random prevalence the variances are indistinguishable from the mean and for fixed prevalence the variances are indistinguishable from the x-axis. For any readers interested in the contribution of the random prevalence to the variance in number of kin, the two variances are shown in Figure B-1. Recognizing the stochastic nature of receiving a cancer diagnosis increases the variance by orders of magnitude.
- The agreement of variance and mean for the random prevalence model is compatible with a Poisson distribution of the age-standardized numbers of kin. Because the numbers are small, the mean number is also a good approximation to the probability of having at least one kin with the condition.
- The age-standardized prevalences are small, because cancer incidence in Japan is also low (see the incidence rates in Figure 2, which reach maxima of only about 0.015 in 1975 and 0.025 in 2018.
- There are marked differences between 1947 and 2019 in Focal’s experience of relatives being diagnosed with cancer.
  – The values for children are slightly lower in 1947 than in 2019. The values for grandchildren and great-grandchildren are higher under 1947 rates than under 2019 rates, reflecting the growth of descendants in the former case relative to the latter.
  – For ancestors, the numbers of kin are larger under 2019 rates, reflecting the greater longevity of parents, grandparents, and great-grandparents under those rates.
  – The differences between years for siblings, nieces-nephews, aunts-uncles, and cousins are smaller, and sometimes favoring one year, sometimes the other.

### 5.3 Dependency ratios

Dependency ratios quantify the relative abundance of subsets of a population that are thought to have different functions relative to each other. They have the form

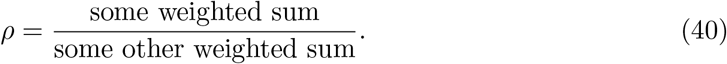

The usual economic dependency ratio uses sums of individuals in age ranges considered to be ‘dependent’ (0–16, 66-*ω*) and independent or ‘productive’ (16–65). Sometimes the young and old dependency ratios are reported separately, because young and old dependent individuals tend to receive support from different sources (from parents and from the labor force, respectively). Loichinger et al. (2017) and Prskawetz and Sambt (2014) discuss a variety of extensions of the idea, taking into account actual economic activities rather than treating everyone within a fixed age interval as equivalent.

The kinship model makes it possible to calculate dependency ratios within parts of the kinship network. It may be that dependencies among her relatives are of more immediate interest to Focal than are dependencies among the whole population. The stochastic version of the kinship model provides variances in the dependency ratios of kin, as a function of the age of Focal.

Consider the traditional age-defined by age limits. Define an output vector **y** as a linear transformation of the vector **k** given by a matrix **W**, of size 3 × *ω*. The first row of **W** contains ones for young dependent ages, the second row contains ones for productive ages, and the third row contains ones for old dependent ages. All other entries are 0. Equation (32) is a small example.

The vector 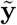 containing the means and covariances of the three age groups is given by

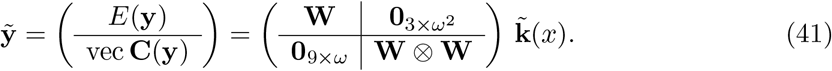

The young and old dependency ratios are

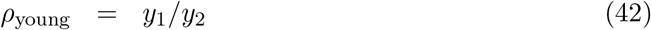

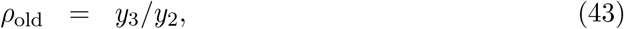

respectively. The dependency ratios are ratios of random variables. The means and variances of ratios of random variables are not given by the ratios of the corresponding means and variances. Instead, the means and variances of the young dependency ratio are given, to second order, by

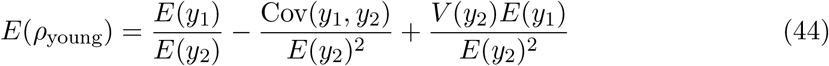

and

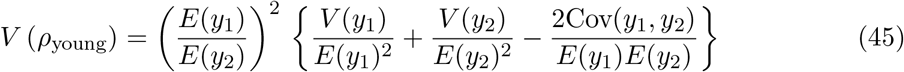

(Johnson, Kotz, and Kemp 1993, p. 55, Stuart and Ord 1987, Sec. 10.5). The corresponding expressions for the mean and variance of *ρ*_old_ are obtained by substituting *y*_3_ for *y*_1_ in equations (44) and (45). All the necessary means, variances, and covariances of the *y*_*i*_ are contained in the vector 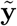 calculated in equation (41).

There is a technical issue in calculating ratios if there is a non-zero probability of the denominator, *y*_2_ in this case, being zero, in which case the ratio is undefined. See Appendix A.3.

#### 5.3.1 Prediction intervals for dependency ratios

To calculate prediction intervals, we need to impose a probability distribution on the dependency ratio. The ratio is continuous and non-negative, so the Poisson distribution that applied to kin numbers is not appropriate. The gamma distribution, which has support on the non-negative real line, is flexible in shape, and includes the exponential, Weibull, χ^2^, and Erlang distributions as special cases, is an attractive choice. For a variable *ξ*, the shape and scale parameters *a* and *b* of the Gamma distribution can be estimated by the method of moments as

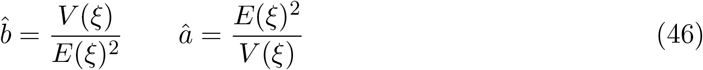

The desired prediction intervals are then obtained from the inverse of the Gamma distribution.

#### 5.3.2 An example: Kinship dependency ratios for Japan

For what types of kin should we calculate dependency ratios? There are many options, some less informative than others. Dependency calculations are perhaps of limited interest for individual kin types, which may only span a limited age range (e.g., Focal is unlikely to have both young and old dependent children, will never have young dependent grandparents, and so on). However, for larger portions of the kinship network, dependency can reveal something about the age patterns among the kin, in a broad sense, of Focal.

In Figure 7 we show the dependency ratios and their prediction intervals for Focal’s entire kinship network as shown in Figure 1 and also for a set of kin comprising a closer family (parents, grandparents, siblings, children, and grandchildren).

**Figure 7:**
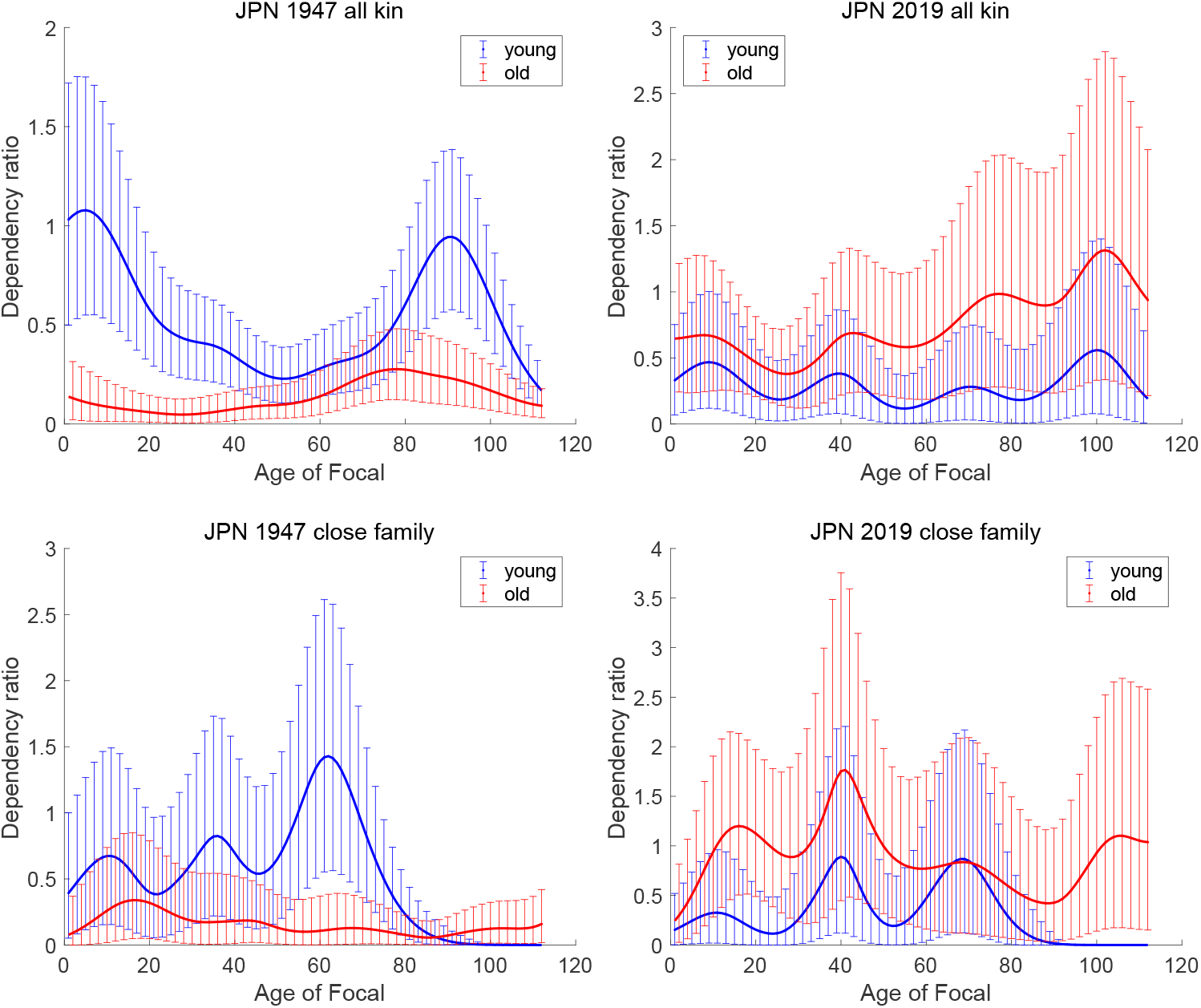
The young and old dependency ratios among all kin of Focal and among immediate family (parents, grandparents, siblings, children, grandchildren) as a function of the age of Focal. The error bars are 90% gamma distribution prediction intervals. For clarity error bars are plotted for every other point.

- The dependency ratios among kin are large, reaching values as large as 2–6; that is, 2–6 dependent kin per each productive kin in the family. This suggests interesting differences between kinship networks and populations as a whole in terms of production and dependence.
- Under 1947 rates, the mean young dependency ratio is consistently greater than the old dependency ratio, because the high fertility in 1947 means that there are many dependent children, grandchildren, and great-grandchildren.
- Under 2019 rates, the pattern is reversed. The old age dependency ratio is consistently greater than the young age ratio. This is a reflection at the kinship network level of the aging of the population under 2019 rates.
- The prediction intervals for dependency ratios are wider under 2019 rates than under 1947 rates, especially for close family kin. The numbers of kin needing support are less certain in the smaller family situation in 2019.

## 6 Discussion

The stochastic kinship model can be written in concise form as combining a state equation and an output equation; when the output is a linear transformation of the kin vector, then the model is

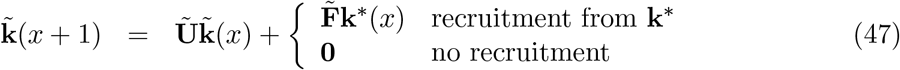

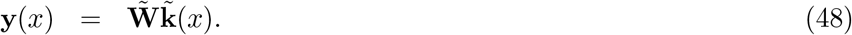

Comparing equation (47) with (1) shows that the stochastic model can follows the procedure for the deterministic model closely, given the right matrices. A protocol for calculating the matrices and carrying out the analysis is given here.

### 6.1 A protocol for calculating a stochastic kinship network

The following is a step-by-step protocol for constructing and analyzing the stochastic kinship model. It assumes as a starting point the time-invariant, age-classified, one-sex model, with the androgynous approximation providing two-sex results. For extensions that relax these conditions, see Section **??**

1. Obtain age-specific schedules of mortality and fertility. Use these to construct the survival matrices **U** and **F**.
2. Obtain or create a distribution π of ages at maternity. In the absence of other information, π can be calculated from the fertility schedule and the stable age distribution as in Caswell (2019, Eq. 3).
3. Construct the stochastic projection matrices Ũ and 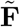 as in equation (4) with the sub-matrices **D**_U_ and **D**_F_ as given by equations (8) and (9).
4. If desired, construct the matrix 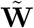 to calculate an output as a linear transformation of the kin vector. This paper has shown several examples of outputs 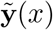 but these by no means exhaust the possibilities.
  a. To compute numbers or weighted numbers of kin, along with their variances and standard deviations over any desired age range, the transformation matrix **W** is described in Section 3.1 and the output matrix 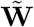 is given in equation (28).
  b. To calculate age-standardized numbers of kin exhibiting some condition, based on age-specific prevalence rates, use the output matrix 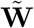 in equation (36), with the matrix **D**_*w*_ = **0** for fixed prevalences or as give by equation (38) for random prevalences.
5. Specify the initial condition for Focal as in equation (10).
6. Specify the initial condition for the mother of Focal as the distribution of a single individual drawn from the π, as in equation (13).
7. Proceed through the kin types in Figure 1 in alphabetical order.
  a. For each kin type, specify the initial condition 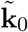 as either **0** or as a mixture, over the distribution of age at maternity, of some other kin type. Follow the procedure in Section 2.4, Item 4, applying equation (18) to each of the mixtures specified in Table 2.
  b. Iterate the state and output equations (47) and (48) to project the vector 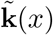 and output vector 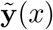 using Ũ, 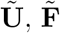, and 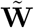 from age *x* = 0 up to a desired maximum age of Focal.
  c. If desired, obtain two-sex results under an androgynous approximation by applying the GKP factors to 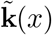 as in Section 3.2.
  The result is a set of kin population vectors 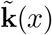 containing the means of, and covariances among, all age classes, for all the types of kin included. These permit other calculations, such as:
8. Calculate prediction intervals for kin numbers using binomial or Poisson distributions as in Section 4.2.
9. Use the Poisson or binomial cumulative distribution functions to calculate the probability of having at least any specified number (one, two, etc.) of kin, as in Section 5.1.
10. Calculate ratio statistics (e.g., dependency ratios) based on selected ages using the results for means and variances of ratios of random variables in Section 5.3.

### 6.2 Implementation: Big state spaces and big matrices

The matrices 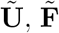, and 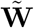 project the covariances as well as the means of the kin structure.

The resulting matrices are large. It is advisable to implement the model using a programming language with sparse matrix capabilities (e.g., Matlab, Python, R). This makes it possible to store the large matrices involved. In the example presented here, with 112 age classes, Ũ and 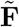 have 112 + 112^2^ = 12, 656 rows and columns, but are very sparse. Sparse matrix calculations also greatly increase the speed of the calculations. The stochastic calculation of the complete kinship network shown in Figure 1 requires only about 7 seconds in Matlab on a very ordinary laptop computer.

It is conceptually straightforward to extend the stochastic model to include additional demographic processes, just as the deterministic model has been extended. Some of these extensions will lead to very large matrices.

- Time-variation in the demographic rates could be included following the approach in Caswell and Song (2021); the matrices **U** and **F** would replaced by time sequences of matrices, **U**_*t*_ and **F**_*t*_. The matrices Ũ and 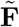 would have the same form used here, but their entries would change over time.
- A two-sex model (Caswell, 2022) would have 112 age classes each for males and for females; **U** would thus have 224 rows and columns, and **U** ⊗ **U** and **F** ⊗ **F** would each have 50,176 rows and columns.
- Multistate models (Caswell, 2020), with individuals classified jointly by age and some other factor (e.g., parity) would have an even larger individual state spaces, and the matrices would be correspondingly larger. For example, with 112 age classes and 6 parity states, **U** and **F** would have 672 rows and columns, and **U** ⊗ **U** and **F** ⊗ **F** would each have 451,584 rows and columns.
- Models for bereavement due to loss of kin include causes of death by augmenting **U** with a block containing transitions from living states to dead states defined by all combinations of age at death and cause of death Caswell, Verdery, and Margolis (2023). The size of the resulting Kronecker products would depend on how many causes of death are included. If 10 causes of death were analyzed over 112 age classes; **U** would have 1232 rows and columns, and **U** ⊗ **U** and **F** ⊗ **F** would have over 1.5 million rows and columns. Big chunks of these matrices would be zero, especially the parts that represent the transition from dead to living stages and recruitment of dead individuals.
- The model presented here provides only the first (mean) and second (covariance) moments. Pollard (1966) and Tuljapurkar (1982) considered higher moments in some related stochastic models. To calculate the third moments would require a matrix in-cluding **U, U** ⊗ **U**, and **U** ⊗ **U** ⊗ **U**. The matrices Ũand 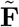would each have more than 1.4 million rows and columns.

The large size of these matrices reflects the extra information incorporated into the model by including stages, time variation, sexes, causes of death, and so on, and the way that these factors influence both means and variances. The good news is that the model extends readily to give stochastic results for all those types of models. And, while the numbers are impressive, there may be no reason to be afraid of large matrices (e.g., Langville and Meyer, 2006).

### 6.3 Demographic stochasticity is different from parameter uncertainty

The demographic stochasticity in this model arises because survival and reproduction are random processes. The model, like all demographic models, applies survival and fertility rates to the individuals in the population. Realizations of the population will differ among themselves because of chance outcomes of the survival and fertility, the rates of which are constant.

That variance among realizations is the result of a given set of demographic rates, specified exactly. If those rates are estimated from data, their exact values will be uncertain. To quantify the variance resulting from that uncertainty requires Monte Carlo sampling from the distribution of the parameter estimates and running the model for each sampled set of parameters. For example, the current United Nations world population projections provide posterior distributions of mortality and fertility using Bayesian methods. Alburez-Gutierrez, Williams, and Caswell (2023) sampled these distributions to generate (deterministic) projections of kinship of all countries, surrounded by uncertainty bounds. The same could be done with the stochastic model; the result would reveal the uncertainty in the variance resulting from the stochasticity.

### 6.4 Independence of individuals and the role of microsimulations

The independence of individuals has been invoked repeatedly in this analysis. That independence is fundamental to the branching process formulation of population growth (Harris, 1963). Much of formal demography is based, explicitly or implicitly, on branching processes. The population projection matrix is the expected value operator of a multitype branching process, the life table and the intrinsic rate of increase are expectations of age-dependent branching processes. All these calculations assume independence of individuals.

When individuals interact, their outcomes are no longer independent, and the age distribution, or the age-stage distribution, or the age-sex distribution, is no longer a valid state variable (a so-called p-state) for the population (Metz and Diekmann, 1986; Metz and de Roos, 1992; Caswell and John, 1992). Instead, the population state variable must account for individuals and the configurations that describe the interactions (Caswell and John, 1992). This is what is done in microsimulation models (e.g., Zagheni, 2015), and for these cases microsimulations are not only possible but essential. Important cases of interactions include pairing of individuals into marriages or partnerships, affinal kin resulting from such pairings, and interactions that take place among members of households.

Microsimulations follow stochastic trajectories of individuals through state space, and thus can provide information on variances as well as means. The branching process model presented here does so as well, without requiring simulations and using the same matrix projection framework as the deterministic model, but, again, only when interactions are not important.

In the context of kinship models, there are a number of interesting examples of microsimulation approaches, using the program Socsim. To cite a few examples: Wachter (1997) analyzed the availability of kin as resources for the elderly, and how that is likely to change in the future. Murphy (2011) studied the effects of the demographic transition in Britain on spouses, children, grandchildren, parents, grandparents, and siblings. Zagheni (2011) used microsimulation to study the impact of HIV/AIDS on the frequency of orphans and double orphans, and the availability of extended kin (grandparents, aunts, uncles, siblings) as resources for those orphans. Verdery and Margolis (2017) used Socsim to examine the prevalence of kinlessness, including partners, children, siblings, and parents. Verdery et al. (2020) studied the impact of COVID-19 on the loss of kin using a bereavement multiplier applied to grandparents, parents, siblings, spouses, and children. Alburez-Gutierrez, Mason, and Zagheni (2021) used microsimulation to study “sandwich” generations: individuals confronting care demands from young children and elderly parents. Snyder et al. (2022) examined the impact of COVID-19 on bereavement due to loss of grandparents, parents, siblings, and children. This list could be extended.

These studies have, for good reason, focused on temporal changes, demographic events such as pandemics, or differences between population subgroups. They have mostly been restricted to subsets of kin that are of particular interest in those contexts. It is notable that most of the studies examine, if not actual individual interactions, at least relationships between different kin types. Some interactions between kin types may be addressed by the matrix kinship model or extensions of it (e.g., Croll and Caswell, 2024). Others are challenging open research problems.

### 6.5 In conclusion

It is now possible to include demographic stochasticity into the matrix kinship model and to obtain not only means but covariances of the age distributions of kin. The stochastic model is a minimal extension of the deterministic model, and follows the same mathematical framework, but with matrices and vectors expanded to incorporate the stochastic aspects of kinship dynamics. The formulation is shown here for a time-invariant, age-classified, one-sex model, but those restrictions can be relaxed at the cost of large matrices. The computational resources needed for those extensions is an open research problem.

The example calculation of kinship networks for Japan compares a population before and after a demographic transition. Jiang et al. (2023) have compared demographic transition effects on kinship networks in China, India, and Japan, using the deterministic model. Feng, Song, and Caswell (2024), also using a deterministic model, have looked at racial disparities in dementia burden in the United States. Other examples exist. It will be interesting to include variances along with means in such comparative studies. Particularly exciting will be comparisons of the patterns of variance implied by a set of demographic rates with those measured in high-resolution register data, such as the analysis by Kolk et al. (2023) of the Swedish ‘kinship universe.’

## 7 Figures

### 7.1 Means and variances

### 7.2 Means and prediction intervals

### 7.3 Fraction with at least one kin

### 7.4 Prevalence of cancer diagnosis

### 7.5 Dependency ratios and prediction intervals

## 8. Acknowledgements

This research was supported by the European Research Council under the European Union’s Horizon 2020 research and innovation program, ERC Advanced Grant 788195 (FORMKIN). I thank those who have discussed and collaborated on this project, and also the many people who have asked “what about variance?” at every presentation of the kinship model.

## A Useful covariance results

Collected for the convenience of the reader.

### A.1 Covariance of linear transformations

Let **x** be a random vector with mean *E*(**x**) and covariance matrix

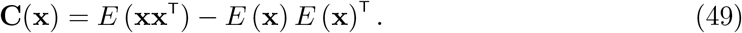

Suppose that **y** is a linear transformation of **x**, given by

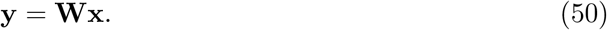

Then the mean of **y** is the linear transformation

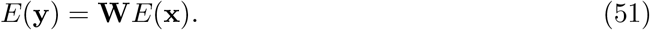

The covariance matrix of **y** is

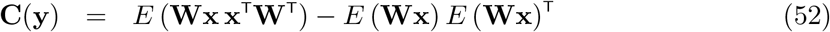

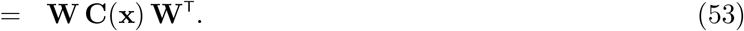

Applying the vec operator to this quadratic form gives

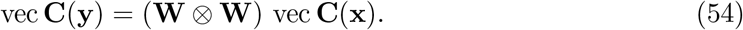

Expressions containing this Kronecker product appear repeatedly in the stochastic kinship model. The projection of the kin vector 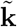 is a sum of two linear transformations, with matrices **U** and **F**, leading to the Kronecker products in equation (4). Calculations of sums and weighted sums of the kin age structure similarly create an output vector as a linear transformation of the kinship vector, leading to the Kronecker product formulation of equation (28).

### A.2 Total variance and total covariance

A result often called the law of total variance says that, if *ξ* is a scalar random variable distributed according to a mixture distribution πwith means *m*_*i*_ and variances *v*_*i*_ conditional on group *i*, then

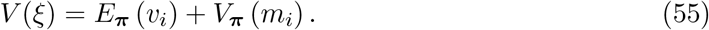

The first term is the within-group variance and the second term is the between-group variance.

The law of total covariance is similar. Let ***ξ*** be a vector-valued random variable distributed according to a mixture distribution **π**, with means *E*(***ξ***_*i*_) and covariance matrices **C**(***ξ***_*i*_), again conditional on group *i*. Then

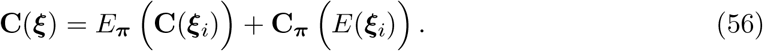

The first term is the within-group covariance and the second term is the between-group covariance.

### A.3 Means and variances of products and ratios

Results on the moments of products and ratios of random variables are scattered. It is useful to collect some of them here. They depend on whether or not the variables can be considered independent.

If *x* and *y* are independent scalar random variables, then the mean and variance of their product are given by

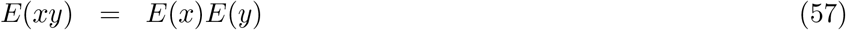

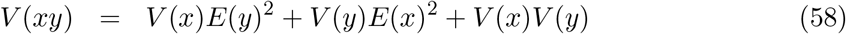

(Stuart and Ord, 1987; Goodman, 1960).

If **x** and **y** are independent vector random of the same length, the mean and variance of their scalar product are

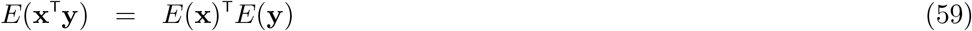

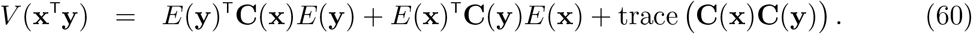

For derivation, see https://stats.stackexchange.com/a/405723. The terms in equation (60) correspond to those in equation (58).

If *x* and *y* are not independent, then

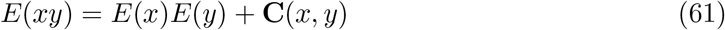

and

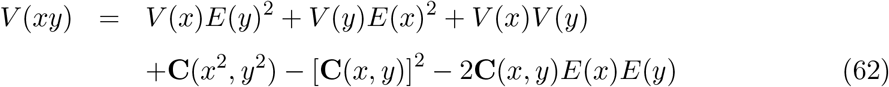

(Frishman, 1971, Sec. 2).

The expectation of the ratio of two random variables, not necessarily independent, is obtained by Taylor series expansion, resulting in

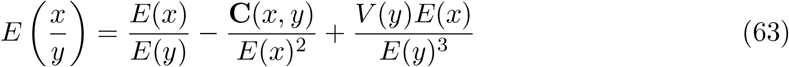

(Johnson, Kotz, and Kemp, 1993, p. 55).

The variance of the ratio of *x* and *y*, again not necessarily independent, is

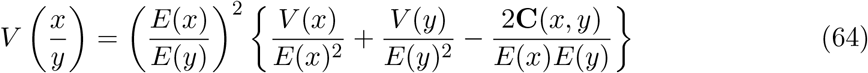

(Stuart and Ord (1987, vol. 1, Sec. 10.5) and Johnson, Kotz, and Kemp (1993, p. 55)).

The analysis of ratios requires some care when the variable in the denominator may be zero. One solution is to replace the distribution of *y* with its zero-truncated distribution. In the context of the dependency ratio in Section 5.3 this would correspond to agreeing not to calculate the kin dependency ratio for an individual with no non-dependent kin. Given the Poisson distribution of kin numbers, this would use a zero-truncated Poisson distribution (Patil et al., 1984).

The zero-truncated Poisson distribution with mean *λ* has a probability mass function

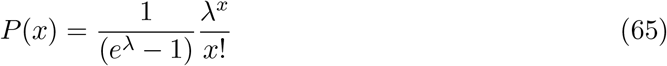

with mean and variance

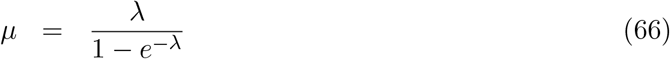

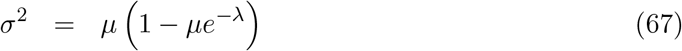

(Patil et al., 1984, p.28). These results could be applied directly to equations (44) and (45). The zero-truncation correction for the covariance terms is an open problem.

## B Variance in cancer prevalence: fixed vs. random

**Figure B-1:**
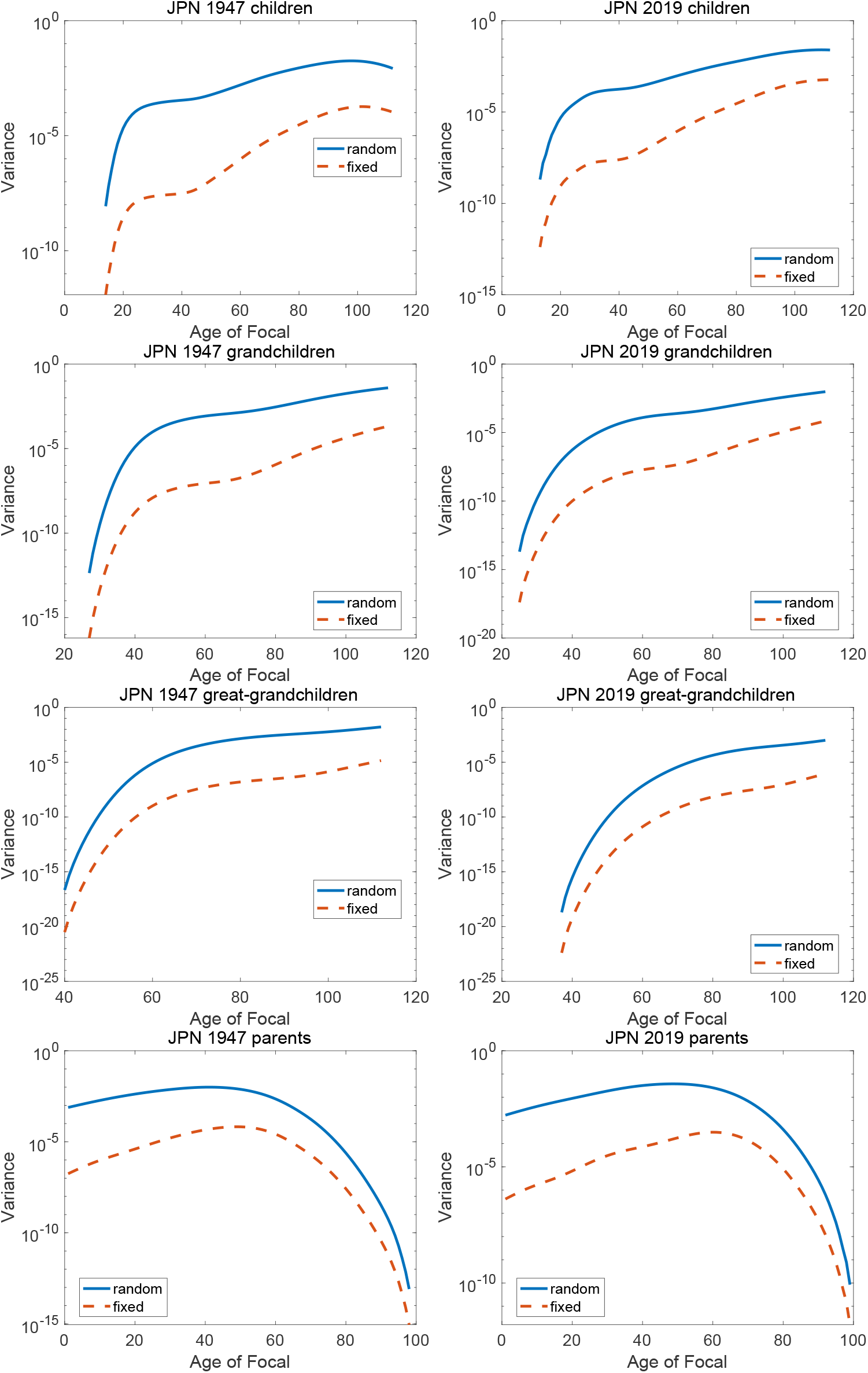
(part 1) Variance in the number of kin experiencing a cancer diagnosis (‘incidence’) as function of the age of Focal, calculated assuming fixed and random prevalences of cancer. Note the differing scales on the y-axis.

**Figure B-1:**
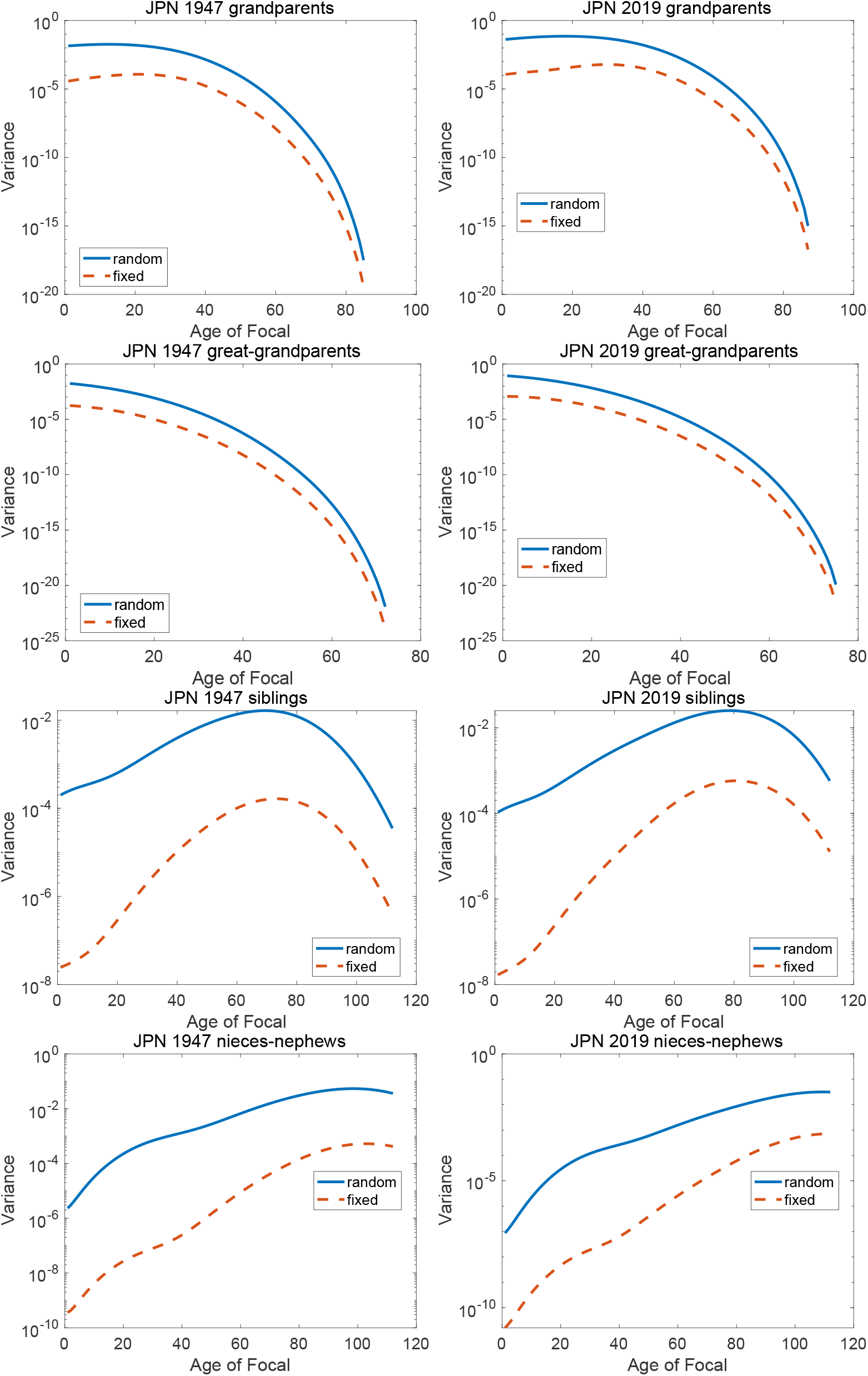
(part 2) Variance in the number of kin experiencing a cancer diagnosis (‘incidence’) as function of the age of Focal, calculated assuming fixed and random prevalences of cancer. Note the differing scales on the y-axis.

**Figure B-1:**
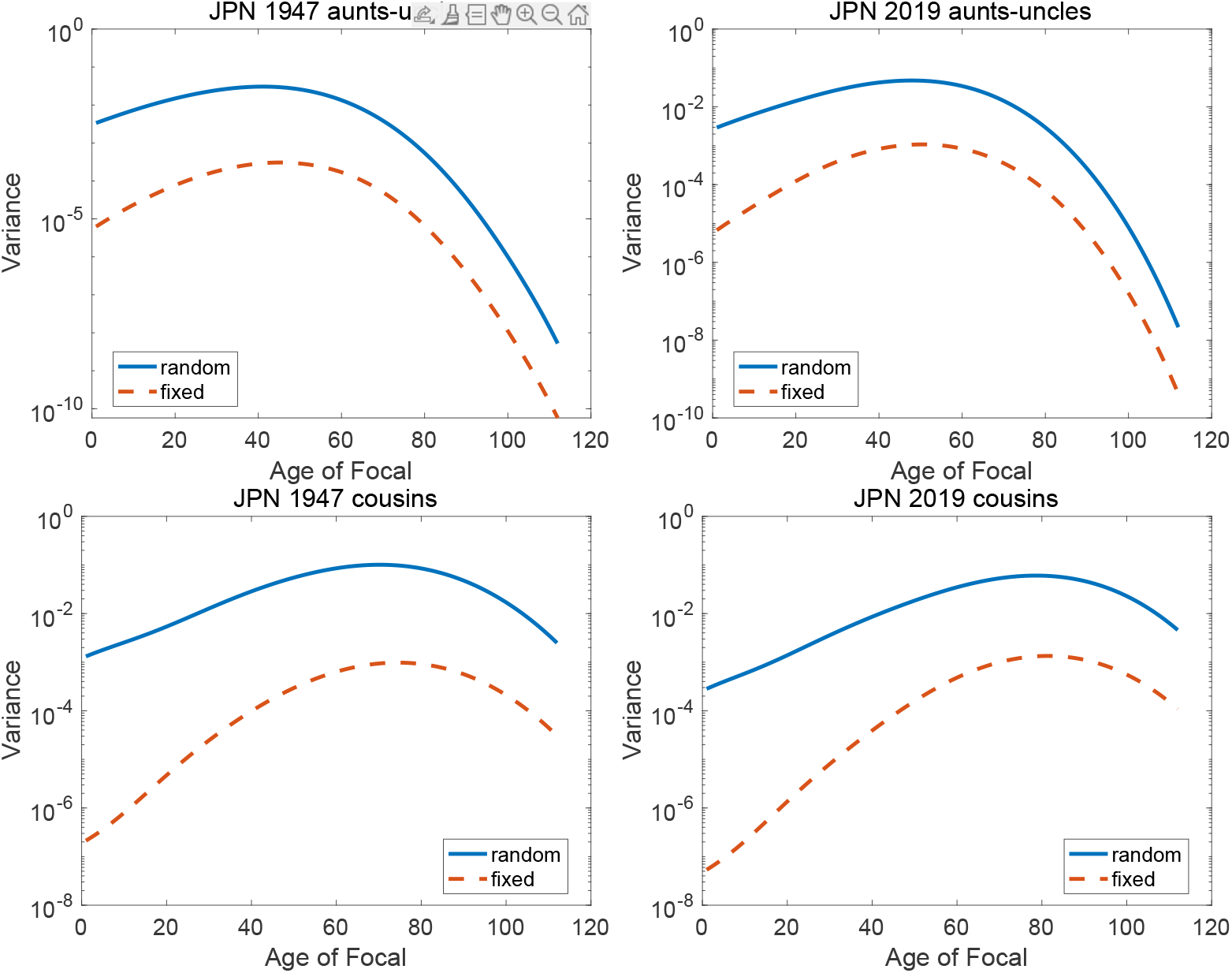
(part 3) Variance in the number of kin experiencing a cancer diagnosis (‘incidence’) as function of the age of Focal, calculated assuming fixed and random prevalences of cancer. Note the differing scales on the y-axis.

## Footnotes

1 Stochasticity enters into demographic analysis at three levels: individual stochasticity, demographic stochasticity, and environmental stochasticity (Caswell, 2001, 2009). Individual stochasticity is the random outcome of probabilistic events in the life of an individual. Two individuals subject to identical demographic rates will differ in their lives; the distribution of longevity implied by a life table is a familiar example. The canonical model for individual stochasticity is an absorbing Markov chain. Demographic stochasticity leads to random differences among populations when individual stochasticity is combined with a renewal process that creates new individuals. Two populations subject to identical demographic rates will diverge in size and structure due to chance outcomes of survival and reproduction. The canonical model is a branching process. Environmental stochasticity is produced by random changes in the conditions in which individuals live, thus changing the demographic rates to which individuals are subject. Two populations in environments characterized by identical stochastic processes will diverge in size and structure because of random differences in the demographic rates to which individuals are subject. The canonical model for environmental stochasticity is a product of random matrices (e.g., Cohen, 1976; Tuljapurkar, 1982, 1990).

2 This is said with tongue firmly in cheek. Certainly for this author, and probably for most of the readers of this paper, these results will not be a reminder. That’s OK.

3 The Kronecker product formulation in equation (4) has appeared, independently, in a variety of contexts. Harris (1963) presented the covariance of a multitype branching process but had not applied the vec operator that would have produced the Kronecker product formulation. That was done by Pollard (1966) as a stochastic extension of the deterministic Leslie matrix model. Conlisk (1969) arrived at the expression as a solution for the equilibrium covariance matrix of a vector-valued autoregressive time series model. Feichtinger (1971) presented a detailed derivation of Pollard’s (1966) model in the context of stochastic demography. Co-hen (1977) presented a Kronecker product formulation for the first and second moments of population in a stochastic environment. Tuljapurkar (1982) (see also Tuljapurkar 1990, Chap. 7) used results from stochastic dynamical system theory (Bharucha, 1961) to obtain the moments in correlated, Markovian, stochastic environments in terms of Kronecker products of projection matrices. Cohen and Tuljapurkar both derived the asymptotic growth rates of the moments under environmental stochasticity. Bartholomew (1982) arrived at the Kronecker product formulation in studying covariances in Markov chain models of social mobility. Caswell and Vindenes (2018) used a similar model to explore the relationship between multitype branching processes and the demographic variance in diffusion models. There are no doubt more examples.

4 This system of equations is familiar from linear systems theory, where the canonical discrete-time system is written (e.g., Zadeh and Desoer, 1963; Schwarz and Friedland, 1965) 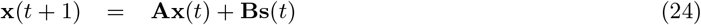 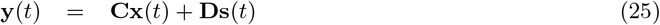 where **x** is the state vector, **s** is the input or stimulus or input vector, **y** is the output vector, and **A, B, C**, and **D** are matrices of appropriate dimensions. The output is a function of the current state and the current stimulus; it does not exhibit the dynamic memory of the state vector **x**, which is updated from its current to its next value.

5 This is not a confidence interval on an estimated mean; the mean is calculated exactly, and is not subject to sampling variation.

6 The value for parents is fixed by the supposition that both of Focal’s parents are alive at birth. In this example, the method of moments estimators of the initial numbers for grandparents and great-grandparents are 4 and 8, respectively, to within thousandths of a percent.

7 These are the presence or absence of beta amyloid A*β* peptides, which are associated with plaque formation in Alzheimer’s disease, the presence or absence of tau protein, which forms filaments in the brain and is also associated with Alzheimer’s, and the presence or absence misfolded alpha-synuclein (*α*-syn) protein, which is associated with Lewy body (LB) dementia.

8 This distinction corresponds to the distinction in survival analysis between the ‘deterministic survival group’ and the ‘random survival group’ (Bowers et al., 1997, Sections 10.3 and 10.4) when interpreting survivorship. It also appears as the distinction between fixed and random rewards in analysis of lifetime fertility using Markov chains with rewards (van Daalen and Caswell, 2015; van Daalen and Caswell, 2017). The fixed reward is equivalent to saying that a fertility rate of, say, 0.3 means that every woman produces exactly 30% of a baby. The random reward interpretation says that every woman of age class *i* has either one baby or zero babies with probabilities 0.3 and 0.7.

9 Data obtained from https://ganjoho.jp/reg_stat/statistics/data/dl/excel/cancer_incidenceNCR(2016-2018)E.xls and https://ganjoho.jp/reg_stat/statistics/data/dl/excel/cancer_incidence(1975-2015)E.xls.

